# Combined promoter-capture Hi-C and Hi-C analysis reveals a fine-tuned regulation of 3D chromatin architecture in colorectal cancer

**DOI:** 10.1101/2022.11.08.515643

**Authors:** Ajay Kumar Saw, Ayush Madhok, Anupam Bhattacharya, Soumyadeep Nandi, Sanjeev Galande

**Affiliations:** Laboratory of chromatin Biology and Epigenetics, Department of Biology, Indian Institute of Science Education and Research, Pune, 411008, India; Division of Life Sciences, Institute of Advanced Study in Science and Technology, Vigyan Path, Paschim Boragaon, Garchuk, Guwahati, Assam, India; Department of Molecular Biology and Biotechnology, Cotton University, Panbazar, Guwahati, Assam, India; Data Sciences and Computational Biology Centre, Amity Institute of Integrative Sciences and Health, Amity University Haryana, Gurugram, Manesar, 122413, Haryana, India; Centre of Excellence in Epigenetics, Department of Life Sciences, Shiv Nadar University, Gautam Buddha Nagar, Greater Noida, Uttar Pradesh, India

**Author notes:** These authors contributed equally to this work (Ayush Madhok, Anupam Bhattacharya). Umeå Plant Science Centre, Department of Plant Physiology, Umeå University, Sweden (Soumyadeep Nandi). To whom correspondence should be addressed. 1. Centre of Excellence in Epigenetics, Department of Life Sciences, Shiv Nadar University, Gautam Buddha Nagar, Greater Noida, Uttar Pradesh, India.;, 2. Data Sciences and Computational Biology Centre, Amity Institute of Integrative Sciences and Health Amity University, Haryana, Gurugram, Manesar, 122413, Haryana, India.

## Abstract

Hi-C is a widely used method for profiling chromosomal interactions in the 3-dimensional context. Due to limitations on the depth of sequencing, the resolution of most Hi-C datasets is often insufficient for scoring fine-scale interactions. We therefore used promoter-capture Hi-C (PCHi-C) data for mapping these subtle interactions. From multiple colorectal cancer (CRC) studies, we combined PCHi-C with Hi-C datasets to understand the dynamics of chromosomal interactions from cis regulatory elements to topologically associated domain (TAD)-level, enabling detection of fine-scale interactions of disease-associated loci within TADs. Our integrated analyses of PCHi-C and Hi-C datasets from CRC cell lines along with histone modification landscape and transcriptome signatures highlight significant genomic structural instability and their association with tumor-suppressive transcriptional programs. Such analyses also yielded nine dysregulated genes. Transcript profiling revealed a dramatic increase in their expression in CRC cell lines as compared to NT2D1 human embryonic carcinoma cells, supporting the predictions of our bioinformatics analysis. We further report increased occupancy of activation associated histone modifications H3K27ac and H3K4me3 at the promoter regions of the targets analyzed. Our study provides deeper insights into the dynamic 3D genome organization in CRC and identification of affected genes which may serve as potential biomarkers for CRC.

**GRAPHICAL ABSTRACT:** 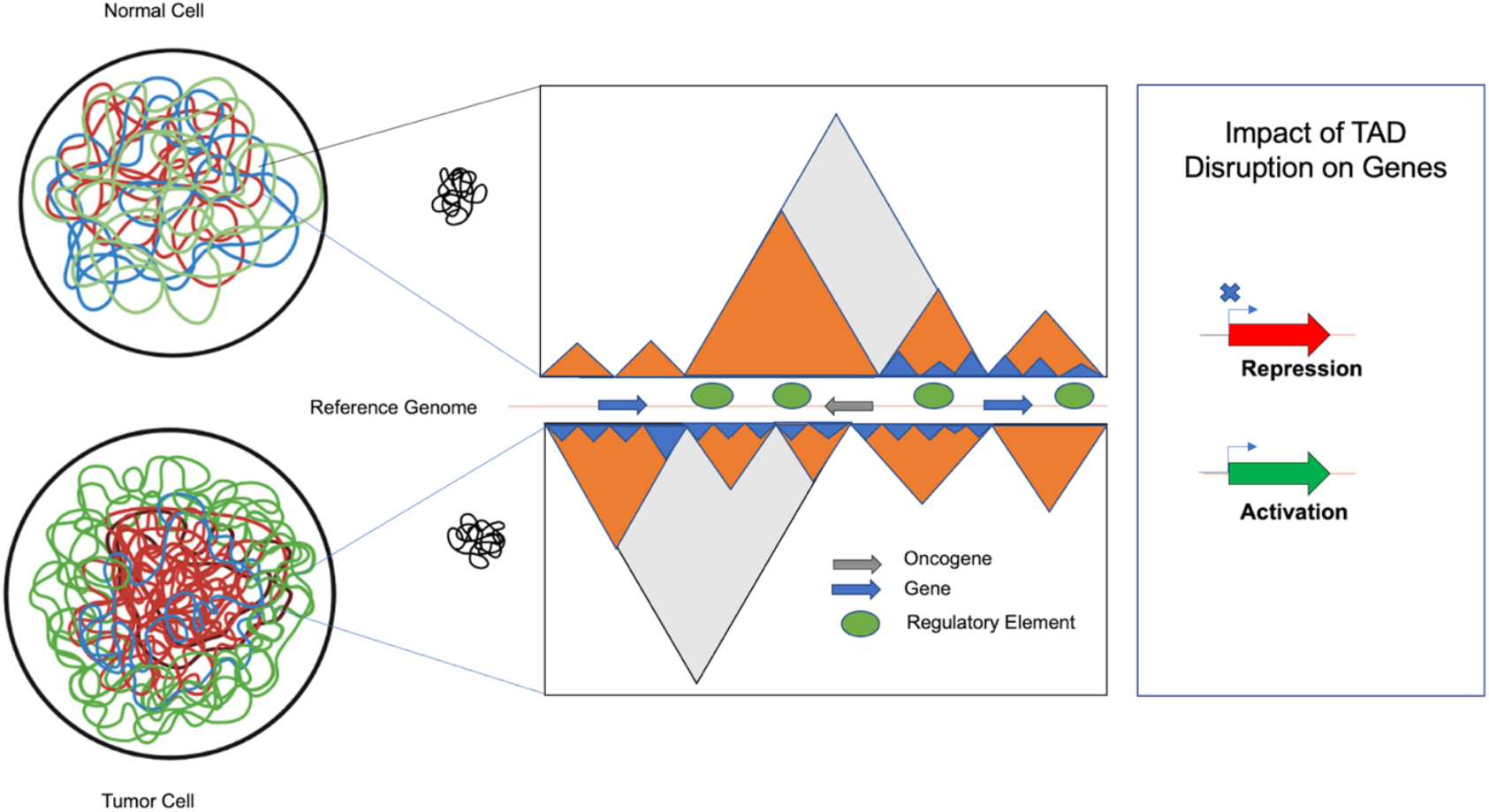

## INTRODUCTION

Until recent past studies related to the impact of three-dimensional changes on gene expression regulation were restricted due to limited tools available to study the 3D genome. Due to the remarkable progress in the development of cutting-edge techniques to decipher chromatin organization, the investigators are exploring the gene expression mechanism in a new dimension. Techniques such as fluorescence in situ hybridization (FISH)^1,2^, and chromosome conformation capture (3C, 4C, 5C, Hi-C, and promoter-capture Hi-C) has been developed for gene expression study. Several studies have taken advantage of these techniques^3–11^, revealing hierarchical layers of spatial organization of genomes^12–21^, and unveiling that 3D conformation of the genome is highly dynamic during animal development. For instance, during the early stage of development genomes exhibit a relaxed chromatin conformation which becomes more compact during later stages. Recent studies have also reported that TADs are much weaker at the early stage of development than in the later stages^22,23^. Thus, it has been postulated that relaxed chromatin conformation during the early stages may facilitate the efficient resetting of the chromatin organization during development. Therefore, it is now evident that the genome conformation is highly dynamic while modulating it’s gene expression profile^24^. Chromosome conformation capture methods such as Hi-C and promoter-capture Hi-C (PCHi-C) provide information about population-averaged ensemble structures, as they are performed on millions of cells. Hi-C maps have been produced in numerous human cell types including embryonic stem cells and early embryonic lineages^25–30^, immune cells^9^, fibroblasts^31^ and other primary tissue types^32^. Hi-C^4^ enables the genome-wide interaction of long-range chromatin contacts and therefore represents a promising approach to identify distal gene targets of disease-associated loci. Hi-C based approaches are at the forefront of guiding our understanding of TAD-level organization, A/B compartments and loop formations^33–41^. For reliable identification of intra-TAD interactions such as the regulatory loops from Hi-C data remains a challenge due to complexity of Hi-C libraries and the substantial cost for the sequencing depth required to achieve statistically significant interactions. On the other hand, targeted chromatin capture techniques such as PCHi-C provide cis-regulatory information for a limited subset of clinically relevant loci at a substantially reduced cost with fine scale chromatin interaction. High resolution maps of the clinically relevant loci allow better prediction of the effects of structural variations and alternation may lead to human disease or developmental abnormalities^42^. PCHi-C greatly increases the capability to detect interactions involving promoter sequences. PCHi-C in distinct cell types recognized thousands of enhancer-promoter interactions and disclosed substantial variations in promoter architecture between cell types and throughout differentiation^43–48^. These studies collectively showed that genome organization reflects cell identity which clearly indicates that disease-relevant cell types are critical for understanding the gene regulatory mechanisms of the disease loci. In support of this concept, several recent studies have employed high-resolution promoter interaction maps to capture tissue-specific target genes of GWAS associations. Javierre et al. produced promoter-capture Hi-C data in 17 primary human blood cell types and captured 2604 potentially important genes for immune and blood-related disorders, including many genes with unannotated roles in those diseases^45^. Mumbach et al. studied GWAS SNPs associated with autoimmune diseases using HiChIP where they captured ∼10,000 promoter-enhancer loops associated with several hundred SNPs to target genes, most of which were not the nearest genes^49^. Montefiori et al. generated high-resolution PCHi-C maps in human induced pluripotent stem cells (iPSCs) and iPSC-derived cardiomyocytes (CMs) where they linked 1999 cardiovascular disease associated SNPs to 347 target genes and demonstrated the importance of considering long-range chromatin interactions when interpreting functional targets of disease loci^50^. Philip et al. conducted a GWAS meta-analysis which contains gene regulatory mechanisms underlying all GWAS risk loci by analyzing PCHi-C to capture chromatin interactions between predisposition loci and target genes, examine gene expression data and integrate these data with chromatin immunoprecipitation-sequencing (ChIP-seq) data for 31 new loci identified together with previously identified loci to the heritable risk of CRC susceptibility^51^.

Colorectal adenocarcinoma, the fourth most common epithelial tumor, continues to be the leading cause of death world-wide^52,53^. Therefore, to gain high-resolution genome architectural insights into colorectal tumor progression, a comprehensive gene regulatory map of human colon cells is required for understanding the impact of TAD disruptions on gene regulation. Here, we present an integrative approach to comprehensively detect structural changes in cancer genomes by combining Hi-C, PCHi-C, RNA-seq, ChIP-seq and single cell RNA-seq (scRNA-seq) analyses for the identification of the multiple target genes which may potentially serve as therapeutic targets for CRC susceptibility (Strategy depicted in Figure 1). We further demonstrate that the expression of these genes as well as associated activatory chromatin modifications such as H3K2754ac and H3K4me3 of the targets are upregulated in CRC. Overall, this study demonstrates the dynamic interplay between global and local chromatin architecture and together integrating with gene expression and chromatin modification profile, we identified novel regions of colorectal cancer susceptibility.

**Figure 1.**
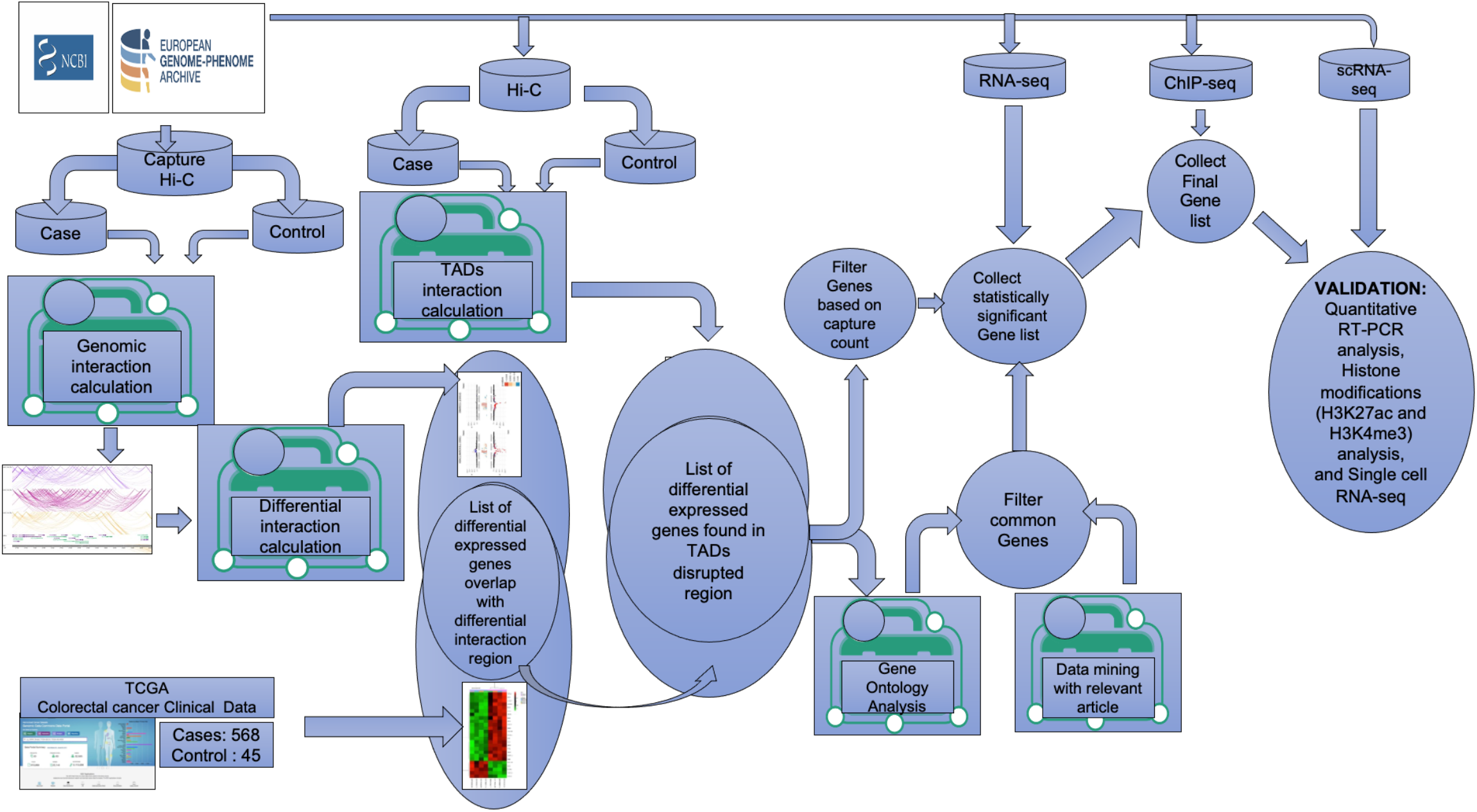
Schematic diagram of the workflow for identification of target genes. Graphical representation of the steps used in the analysis combining multiple large colorectal cancer datasets (PCHi-C, Hi-C, ChIP-seq, RNA-seq and single cell RNA-seq) to identify target genes potentially playing a key role in CRC progression.

## MATERIALS AND METHODS

### Promoter-capture Hi-C (PCHi-C) data processing and analysis

Raw Promoter-capture Hi-C (PCHi-C) sequencing reads were mapped using hg38 human reference genome and filtered using the HiCUP^54^ and bowtie2^55^. For annotation of the promoter interacting regions (PIRs), we used the CHiCAGO tools^56^. Promoter ligated regions (PIRs) are recognized by comparison of the number of promoter-ligated regions in each genomic bin with expected reads according to the generated model. Bins having CHiCAGO scores ≥ 5 are considered to be PIRs. BAM files were converted to a CHiCAGO-compatible format (.chinput) using bam2chicago script. Chinput files are tab-delimited text files which contain basic information of all pairs promoter—”other end” regions for which at least one ligation region was detected. Each line of the chinput file describes one promoter—”other end” pair, this description includes genomic coordinates of the promoter and the “other end”.

### Differential chromosomal interaction analysis using PCHi-C data

Differential signal detection in sequencing data is one of the most common tasks in genomic analyses. We used Chicdiff^57^ to perform differential analysis of significant interactions called by CHiCAGO (between different cell types). This tool combines moderated differential testing for count data using negative binomial generalized linear models implemented in DESeq2^58^ with signal normalization informed by CHiCAGO and nonuniform multiple testing correction.

### Hi-C data processing and analysis

Reads were mapped to the human reference genome hg38 using bowtie2^55^, and then samtools^59^ was used to convert the reads to BAM format. We used Hicexplorer package^60,61^ for Hi-C data processing. “hicBuildMatrix” command builds the matrix of read counts over the bins in the genome, considering the sites around the given restriction site (AAGCTT). Using “hicCorrectMatrix” command, the matrix was corrected to remove GC, open chromatin biases and, most importantly to normalize the number of restriction sites per bin. For topologically associated domains (TADs) calculation, we used “hicFindTADs” command with the following parameters “–minDepth 60000 –maxDepth 120000 –step 20000”. The interaction matrix and other signal tracks were also visualized using “hicplotsTADs” command. TAD boundaries positions were used to analyze localization of detected genomic interactions relative to TAD boundaries.

### ChIP-seq data processing and analysis

ChIP-seq reads were trimmed using the trim galore tool for adapter removal. Filtered reads were mapped to human reference genome hg38 using bowtie2^55^ and then samtools^59^ was used to convert the reads to BAM format. Macs2 callpeak was used for peak calling with --tsize=26, -- gsize= 2.7e9, –extsize 200 and the rest are default settings. MACS2 tools captures the influence of genome complexity to evaluate the significance of enriched ChIP regions and improves the spatial resolution of binding sites through combining the information of both sequencing tag position and orientation.

### RNA-seq data processing and analysis

#### RNA-seq Pipeline 1

The gene expression abundance from pair-end RNA seq datasets were downloaded from the Cancer Genome Atlas (TCGA) data portal. For retrieval of colon cancer datasets, primary sites “colon”, “rectum”, project id “TCGA-COAD”, “TCGA-READ” and sample type “Primary Tumor”, “Tumor”, “Metastatic” were selected. For control datasets, “Blood Derived Normal”, “Solid Tissue Normal” parameter were selected during the data retrieval process. The commands for data retrieval are given below:

##### For case

cases.primary_site in [“Colon”,”Rectum”] and cases.project.program.name in [“TCGA”] and cases.project.project_id in [“TCGA-COAD”,”TCGA-READ”] and files.analysis.workflow_type in [“HTSeq - Counts”] and files.data_format in [“TXT”] and files.data_type in [“Gene Expression Quantification”] and files.experimental_strategy in [“RNA-Seq”] AND cases.samples.sample_type IN [“Primary Tumor”,”Tumor”,”Metastatic”]

##### For control

cases.primary_site in [“Colon”,”Rectum”] and cases.project.program.name in [“TCGA”] and cases.project.project_id in [“TCGA-COAD”,”TCGA-READ”] and files.analysis.workflow_type in [“HTSeq - Counts”] and files.data_format in [“TXT”] and files.data_type in [“Gene Expression Quantification”] and files.experimental_strategy in [“RNA-Seq”] AND cases.samples.sample_type IN [“Blood Derived Normal”,”Solid Tissue Normal”]

The sample manifest file was prepared and GDC client^62^ was used for the downloading of the datasets.

### RNA-seq Pipeline 2

#### Quality control of the dataset and mapping of processed reads

Raw files were downloaded and converted using SRA toolkit. After QC program was used to filter low quality reads and removal of any sequencing errors. The processed reads were aligned by the RNA-Seq aligner STAR(v2.5)^63^ with the Ensemble Human reference annotation (GRCh38).

### Quantifying transcript abundances

RSEM^64^ was used to quantify transcript abundance from the pair-end RNA seq datasets. The human reference genome was built by using the rsem-prepare-reference script. We used STAR to perform transcriptome-based mapping, and gene expression was calculated from STAR-generated BAM files by rsem-calculate-expression scripts.

### Identification of differentially expressed genes

The edgeR package (v3.30.0)^65^, which relies upon count-based expression data for determining differential expression in R was used for differential gene expression analysis. An overdispersed Poisson model was used to account for both biological and technical variability. Empirical Bayes methods were used to moderate the degree of overdispersion across transcripts improving the reliability of inference. Transcripts with zero expression values were removed. Normalization of the counts was performed by using the calcNormFactors function of edgeR, which normalizes for RNA composition by finding a set of scaling factors for the library sizes that minimize the log-fold changes between samples for most genes. TMM method of normalization was used for the datasets.

### RNA-seq Pipeline 3

The preprocessed RNA transcripts were mapped to Homo sapiens (Gencode v29) transcriptomes using salmon^66^. The resulting quant files were imported into R (v4.0.3) for exploratory data analysis. R package tximeta^67^ was used to import and summarize transcript-level data to the gene level. DESeq2^58^ was used for Differential expression analysis with default setting. Further details of used pipeline are available in the following link: (https://bioconductor.org/packages/release/workflows/vignettes/rnaseqGene/inst/doc/rnaseqGene.html).

### GO analysis, network analysis, mutational analysis details

To understand the functional perspective of associating genes with a functional biological term in a systematic way, STRING database was used for the functional annotation of genes. During the analysis, GO biological process libraries 2021, molecular function libraries 2021 was selected. The statistically significant results (p-value <= 0.05) were further considered for downstream analysis as a reference organism for the annotation of genes. The enriched genes clusters were visualized in STRING network. In this study, we primarily emphasized on functional exploration of genes associated in chromatin organization. We curated the function of genes based on functional enrichment analysis associated with key functions like chromatin remodeling, chromatin organization, cellular organization, etc. For mutational analysis: Colon Cancer Atlas and Cosmic database used as reference database having 13,711 CRC tissues and more than 165 CRC cell lines. The affected genes due to TAD boundary changes were mapped against all these CRC cell lines and HT29 and LoVo cell lines.

### Single cell RNA-seq data processing and analysis

We used Seurat package^68^ (v4.0.6) for this integrative multimodal analysis. Genes detected in fewer than 100 cells, cells exhibiting expression of less than 200 detected genes, and cells expressing > 15% mitochondrial genes were removed for downstream analysis. We adopted the general protocol described in Stuart et al.^68^ to group single cells into different cell subsets. We employed the following steps: data normalization, Identification of highly variable features, Scaling the data, Cluster the cells, reorder clusters based on their similarity, run non-linear dimensional reduction (tSNE) and labeling cell types. Principal component analysis (PCA) was carried out on the scaled data of highly variable genes. The first 30 principal components (PCs) were used to cluster the cells and to perform a subtype analysis by nonlinear dimensionality reduction (t-SNE)^69^. Additionally, we utilized SingleR package (v1.8.1) for labelling cell types^70^.

### Cell lines and culture

NT2D1 embryonic cells (Gifted by Dr. Peter Andrews) were cultured in DMEM (Invitrogen), with 20% FBS (Invitrogen), 1X antibiotics (Antibiotic-Antimycotic, Invitrogen) and NEAA supplements (Lonza) at 37° C with 5% CO_2._ The cells were split after every 3 days and maintained at 70% confluency to avoid any differentiation. HT29 colorectal carcinoma cell line (ATCC, HTB-38) was cultured in DMEM, with 10% FBS and antibiotics. Cells were split after every 1.5 days and 2 or 10 million cells were harvested for RNA extraction and chromatin immunoprecipitation assays, respectively.

### RNA extraction and qRT-PCR

The cells harvested (2 × 10^6^), were washed twice with cold 1X PBS and the pellets were resuspended in TriZol (Invitrogen) followed by isolation of total RNA. Following DNase I(Promega) digestion, RNA (260/280 ratio ∼2) was subjected to cDNA synthesis using High Capacity cDNA synthesis kit (Applied Biosystems) as per the manufacturer’s protocol. Target oligos were designed using UCSC genome browser and Primer3 softwares. Expression level of h18s RNA was used as internal control for normalization of target transcripts. Quantitative RT-PCR analyses were performed using TB Green II qPCR master mix (Takara) at the following PCR conditions: step 1, 95°C, 5 min; step 2, 95°C, 45 s, 60°C, 45 s, 72°C, 1 min for 40 cycles using ViiA 7 real-time PCR system (Applied Biosystems). The change in gene expression was calculated using the formula ΔCt = Ct Target − Ct Control. Normalized transcript expression was calculated using the equation 2−(ΔCt), where ΔCt = Ct Target − Ct Control. The oligonucleotide primer sequences used for qRT-PCR analyses are listed in the Supplementary table S1. Statistical analysis was performed using one way ANOVA (Graphpad v9.1) on three biological replicates.

### Chromatin Immunoprecipitation (ChIP) q-PCR

Cells obtained were cross-linked using 1.25% formaldehyde (Sigma) followed by quenching using 150 mM glycine. Cross-linked cells were washed twice with PBS and were subjected to chromatin isolation and shearing as described by Patta et al.^71^ with a few modifications. Briefly, nuclei isolation was performed using the hypotonic buffer (25 mM Tris–HCl pH 7.9, 1.5 mM MgCl_2_, 10 mM KCl, 0.1% NP40, 1 mM DTT, 0.5 mM PMSF and 1× protease inhibitor cocktail (Roche). Pelleted nuclei were lysed using sonication buffer (50 mM Tris–HCl at pH 7.9, 140 mM NaCl, 1 mM EDTA, 1% Triton X-100, 0.1% Sodium deoxycholate, 0.5% SDS, 0.5 mM PMSF and 1× protease inhibitor cocktail (Roche) and chromatin was subjected to the sonication using Covaris M220 sonicator using the following parameters: peak power 75, duty factor 20, burst 300, duration 10 min. We obtained the chromatin fragment size of 200–400 bp. After preclearing the chromatin was subjected to immunoprecipitation using anti-H3k27ac (Abcam), anti-H3K4me3 (Abcam) antibodies overnight at 4°C. Similarly, normal IgG was used as a control. Immunoprecipitated complexes were pulled down by adding Protein A/G beads (Thermo Scientific) and incubated the cocktail at 4°C for 4 h. The immunoprecipitated bead bound chromatin was washed thoroughly using low-salt, high-salt and lithium chloride buffers, and subjected to elution by using the elution buffer (1% SDS, 0.1M NaHCO_3_). The eluted chromatin was de-crosslinked and the protein and RNA contamination was removed by treating with proteinase K (Sigma-Aldrich), and RNase A (Sigma-Aldrich), respectively. Further, the immunoprecipitated chromatin was purified and subjected to the quantitative PCR analysis using the formula ΔCt = Ct _Target_ − Ct _Input_. Fold differences in enrichment were calculated using the equation 2^−(ΔCt)^, where ΔCt = Ct _Target_ − Ct _Input_, for both IgG and TCF1 immunoprecipitated DNA. The primer sequences used for ChIP-PCR analysis are listed in the Supplementary table S2. Statistical analysis was performed using one way ANOVA (Graphpad v9.1) with three biological replicates.

## RESULTS

### Enhancer promoter interactions are enriched in HT29 and LoVo cell lines compared to hESC which correspond to changes in gene expression dynamics

We used PCHi-C data to examine genomic interaction in colorectal cancer (CRC). The captured regions were selected based on genome-wide association studies (GWAS) for Capture Hi-C experiments. The captured regions (genomic region of interest or diseases associated loci) are referred to as “baits” throughout the manuscript. Genome-wide association studies are typically employed to identify genes involved in human diseases. This method searches the genome sequences for small variations called single nucleotide polymorphisms or SNPs that occur more frequently in individuals affected with a particular disease than in the healthy individuals. Law et al. reported a GWAS analysis of 34,627 CRC cases and 71,379 controls of European ancestry that identifies SNPs at 31 new CRC risk loci ^51^. In this study, we selected two colon cancer cell lines (HT29 and LoVo) and compared them with human embryonic stem cells (hESC). We used CHiCAGO^56^ to identify significant interactions. We investigated the interactions that were separated by a distance of at least 10 Kb. We identified bait Interactions with other genomic loci and the results were shown in Figure 2. In total, we identified 1283056 (49%), 1135609 (43%) number of bait interactions in HT29 and LoVo cell lines respectively, and 201356 (8%) interactions in the hESC cell line (Figure 2A). Most interactions were found in cancer cell lines (HT29, LoVo) compared to hESCs. The significant proportions of bait interactions were shared between cancer cell lines compared to hESCs (Figure 2B). We also calculated the counts of interactions with respect to each bait fragment as well as to chromosome-wise across all the cell lines (Figures 2C-D). We observed similar interaction patterns in HT29 and LoVo cell lines compared to hESCs (Figure 2D). We used Chicdiff^57^ for differential bait interaction between cancer versus normal cell lines. We plotted the distribution of differential bait interactions (Figures 3B-C) and observed 3895502 (∼87%) and 582084 (∼13%) differential interactions of HT29 versus hESC and LoVo versus hESC cell lines, respectively (Figure 3A). Differential interactions between two bait fragments (Figure 3D-E) were plotted with interactions within 500 Kb upstream and downstream from bait passing the threshold of 5 (threshold > 5). The threshold criteria >5 was setup by Chicdiff^57^ tool for significant differential interaction. Similar to the differential interaction analysis, we performed differential RNA-seq analysis using CRC datasets obtained from TCGA databases^72^. The identified differentially expressed (DE) genes are depicted in Figure 3F. We found approximately twenty-eight thousand transcripts that were differentially expressed (False Discovery Rate <=0.05). Nearly 62% of transcripts were expressed in cancer cells compared to normal cells. In cancer cells, 35% were protein-coding genes and the rest were non-coding genes (lincRNA, pseudogenes, antisense, miRNA, etc). All the transcripts along with the annotations were provided in Supplementary table S3. Gene expressions related to colorectal cancer (primary tumor, metastatic) were compared with blood-derived normal, solid tissue to determine the DE genes using edgeR^65^. For the extraction of DE genes that lie within the differential interaction region, we overlapped the genomic coordinates of differential bait fragments with the genomic coordinates of DE genes. In hESC/HT29 and hESC/LoVo case studies, we found 24438 and 17385 DE transcripts respectively after overlapping.

**Figure 2.**
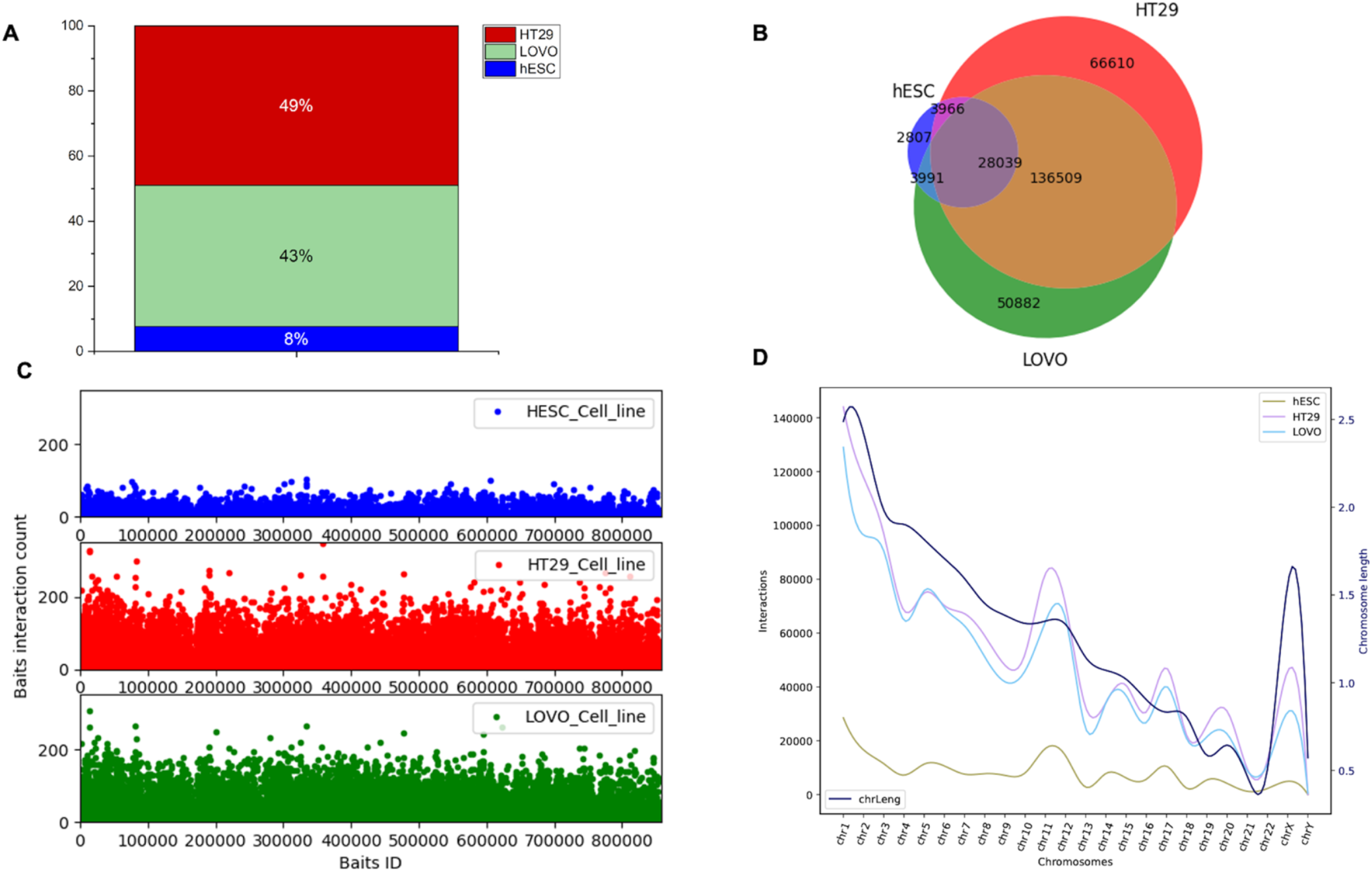
Distribution of promoter capture Hi-C interaction of hESC, HT29 and LoVo cell lines. Here, bait fragment refers to promoter capture region or region of interest and other end fragment refers to those fragments which interact with the bait fragment. (A) Proportion of bait interactions with other ends generated by ChiCAGO^56^ using PCHiC data of HESC, HT29 and LoVo cell lines; (B) Venn diagram displaying the number of bait fragments among hESC, HT29 and LoVo; (C) The interaction counts of bait fragments with other end fragments of hESC, HT29 and LoVo cell lines. (D) Chromosome-wise interaction counts of bait fragments with other end fragments of hESC, HT29 and LoVo cell lines.

**Figure 3.**
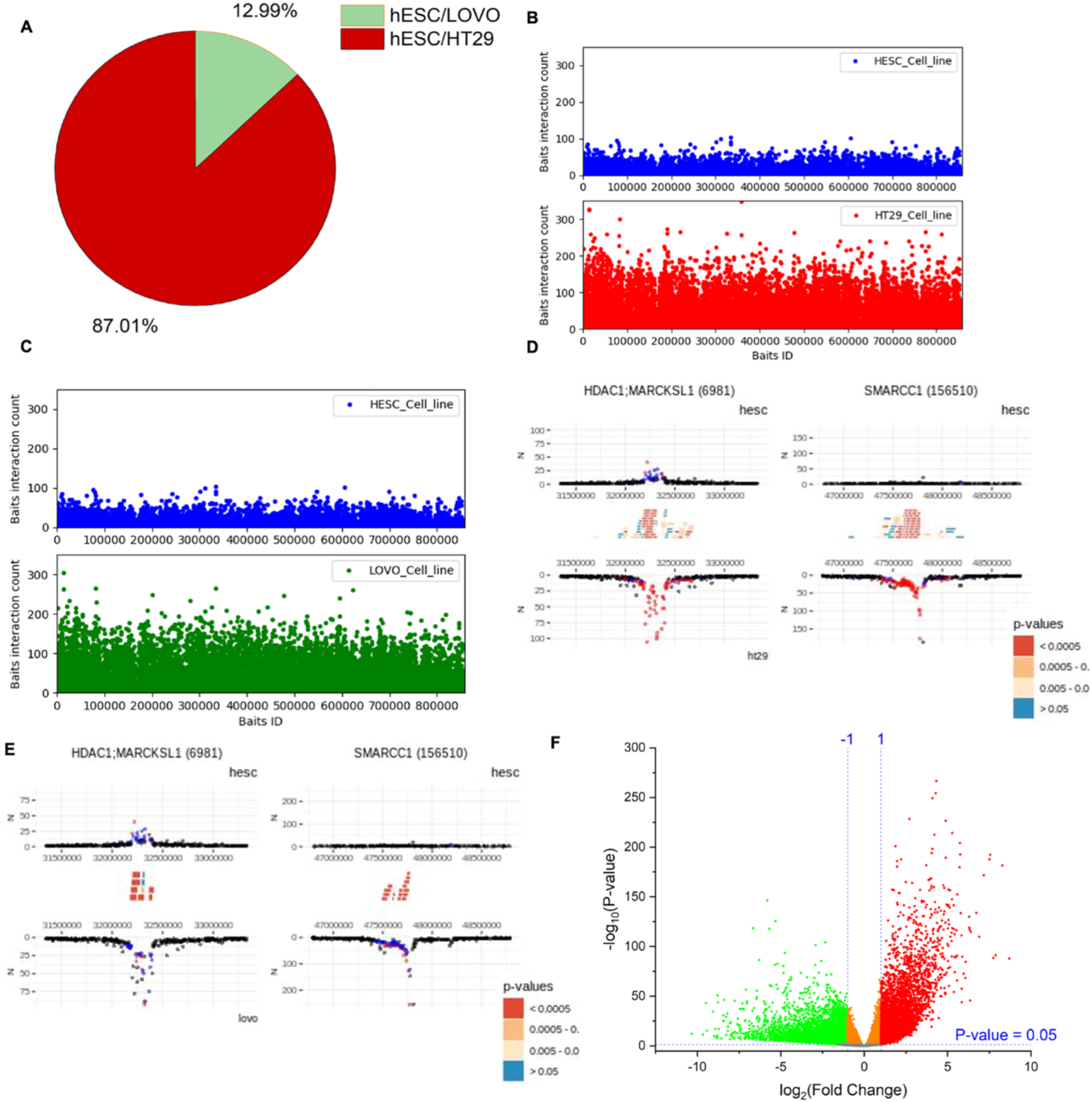
Distribution of differential promoter capture Hi-C interaction between cancer (HT29 and LoVo) versus normal (hESC) cell lines. (A) Proportion of differential bait interactions between cancer versus normal cell lines. (B and C) Differential bait interaction counts of cancer versus normal cell lines with respect to bait fragments. (D and E) Differential interaction detected by Chicdiff^57^ around HDAC1, MARCKSL1 and SMARCC1 genes are plotted. Significant interactions were detected for each condition separately by CHiCAGO^56^, color-coded (blue: 3 < score <=5; red: score > 5). Significant differentially interacting regions detected by Chicdiff^57^ were depicted as red blocks against HT29 versus hESC and LoVo versus hESC cell lines. (F) Volcano plot of differentially expressed genes generated from colorectal cancer dataset from the TCGA database. A total of 57864 transcripts were observed. 27676 statistically significant transcripts were identified (False Discovery Rate <=0.05). 17146 transcripts were expressed in cancer cells while 10530 transcripts were expressed in normal cells.

### Disruption in TAD dynamics of HT29 and LoVo cell lines compared to hESCs involves a subset of colorectal-specific genes

In the previous section, we overlayed differentially expressed genes with the bait fragment’s coordinates of the differential interaction region. To know the distribution of TAD boundary regions with respect to bait fragments, we further overlapped the bait fragments in such a way that the start bait should lie within the range of TAD boundary regions. This helped us to probe the distribution of nearby genes across TAD boundaries. These findings provided insights into better understanding of the role of regulatory mechanisms for structural instability in CRC with respect to normal cells. Using HiCExplorer, we plotted the TADs with 10 Kb resolution and compared the TAD boundaries among the three cell lines (hESC, HT29, and LoVo). We found 8461, 7955 and 6498 TAD boundaries respectively in hESC, HT29 and LoVo cells (Figure 4A). Next, the overlapped TAD boundaries between two cell lines containing at least one DE gene were classified into two categories: conserved TADs and disrupted TADs. The total number of conserved and disrupted TADs (Figure 4B) of hESC/HT29 and hESC/LoVo are 2637 and 5448 and 2138 and 4255, respectively. The criterion for selecting conserved TADs is that the genes contained in the overlapping TADs are the same as genes contained in their corresponding TADs. Hence, overlapping TADs which are not conserved were considered disrupted TADs. We then investigated nearby genes in detail around the TAD disrupted regions. We filtered genes based upon their position in the TAD boundary shift loci between normal and cancer cells (Figure 4C). We mapped and filtered the disrupted genes in TAD shifting boundary regions with the TCGA database (Figure 4D). We found 47% of the disrupted genes were protein-coding and 53% were non-coding genes which include lincRNAs, miRNAs, pseudogenes, and other non-coding variants in the hESC/LoVo cells. Similarly, 40% expressed genes were protein-coding and 60% were non-coding genes in hESC/HT29 cells (Figure 4E).

**Figure 4.**
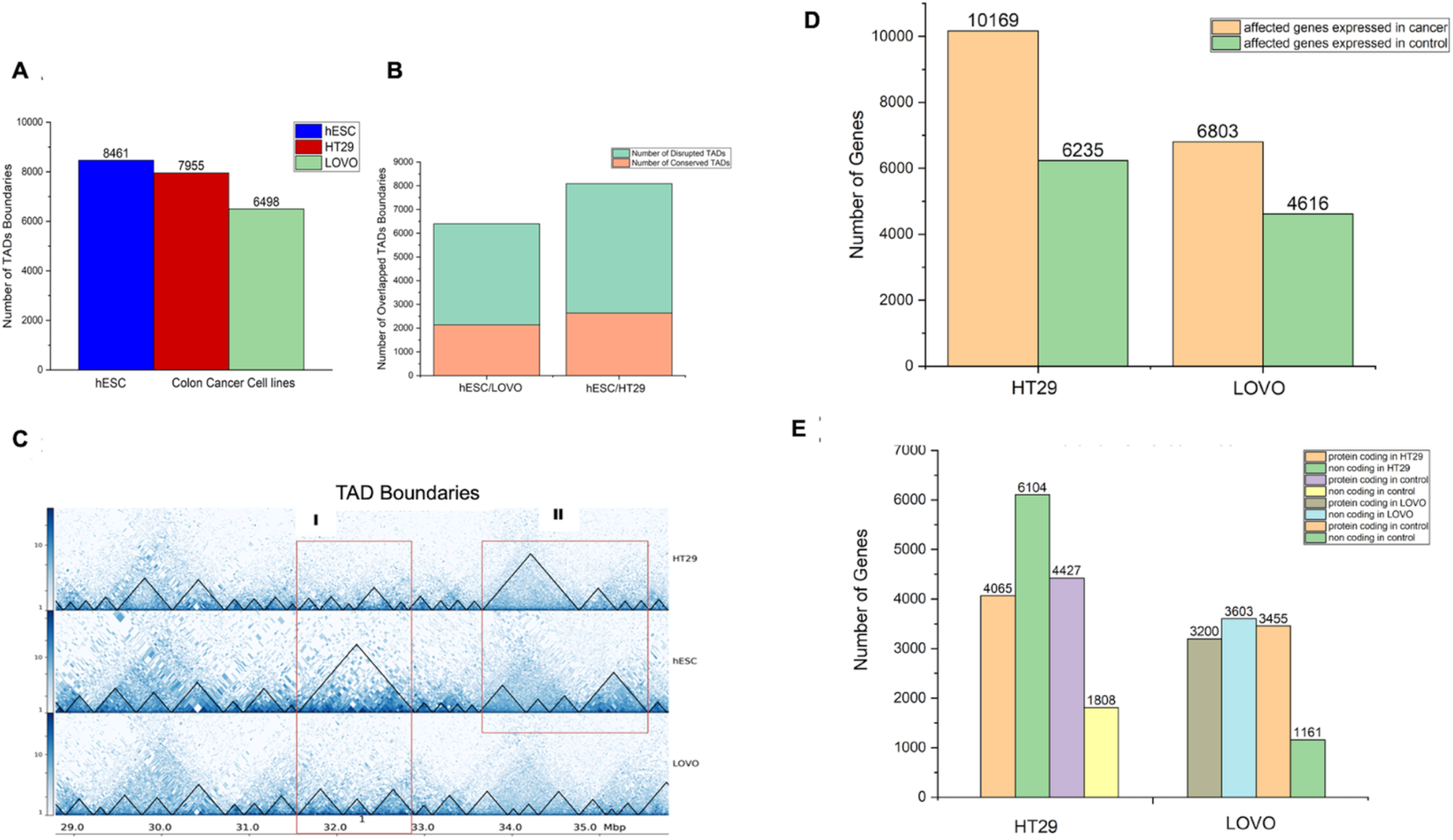
Distribution of genome-wide interactions between CRC (HT29 and LoVo) versus embryonic stem (hESC) cell lines. Here, TAD refers to intra-chromosomal interaction between two different genomic fragments. (A) The number of TAD boundaries for each cell line generated by HiCExplorer^60,61,94^. (B) The number of conserved and disrupted TADs between two cell lines. (C) TAD plot of chr1:28788062-35788062 of HT29, hESC and LoVo cell lines and red rectangle highlight major TAD shift of HT29 and LoVo compared to hESCs. (D) Affected genes due to TAD boundary shifting in HT29 and LoVo while comparing with hESCs. (E) Number of protein-coding and non-coding genes in HT29 and LoVo TADs as compared with hESC TADs.

### Gene ontology, network analysis and mutational study of colorectal-specific genes in the vicinity of disrupted TAD boundaries

A functional analysis revealed that most of the genes found in the disrupted TAD boundaries involved in chromatin remodeling, chromatin assembly and chromatin organization (Table 1). We found SWI/SNF gene clusters, INO80 complexes and histone gene clusters enriched in the datasets. Major protein family that was involved in chromatin remodeling included SMARCB1, SMARCA1, SMARCA2, SMARCA4, SMARCC1, SMARCC2, SMARCA5, SMARCD1, SMARCD2, SMARCD3, ARID1A, ARID1B, INO80E, INO80C, CHD3. Network analysis revealed SWI/SNF regulatory networks, histone family clusters significantly associated with chromatin remodeling, chromatin assembly and chromatin organization (Figure 5A). A gene ontology study also revealed that shifting of TAD boundary affected nearly 300 genes involved in the regulation of transcription factors. Several genes (DMAP1, SIN3B, HIRA, AC002553, NCOR1, SAP30, NCOR2, SIN3A, URI1, AC008985, AC007292, MPND, NCOA6, SAP30L, TRAC) involved in transcription regulation of chromatin binding were affected due to the TAD boundary changes, suggesting the shifting of TAD boundaries affected the gene regulation in CRC. The enriched gene ontology and pathway analyses for both HT29 and LoVo cell lines are provided as Supplementary table S4. Figure 5B depicts a volcano plot showing the list of the affected genes that were involved in chromatin remodeling, chromatin assembly and chromatin organization expressed in cancer cells. The heatmap and normalized count of list of affected genes are listed in Supplementary figure S1. To further explore the changes in genomic instability due to shift in TAD boundaries, we integrated the mutational datasets with the affected genes due to changes in TAD boundaries. We overlapped the affected genes with the CRC database and cosmic database^73,74^ to find if the affected genes were flagged as oncogenes due to the shifting of TAD boundaries. Such analysis revealed that nearly eighteen thousand genes were affected during the TAD boundary alterations in LoVo and HT29 cells compared to the hESCs. Mutational analysis revealed that AR1D1A, ATRX, centromere protein complex, histone complex, chromatin organizers SATB1 and SATB2, SWI1/SNF1 complex and Histone deacetylases were frequently mutated in this dataset. More than 50% of the affected genes due the shifting of TAD boundary harbored mutations in various cancer-specific cells including HT29, LoVo, KM12, LIM1215, LIM1863, SW480, HT115, SNU175, HCC2998, SNUC4, HCT15, CX1, WIDR, CoCM1, etc.

**Table 1:**
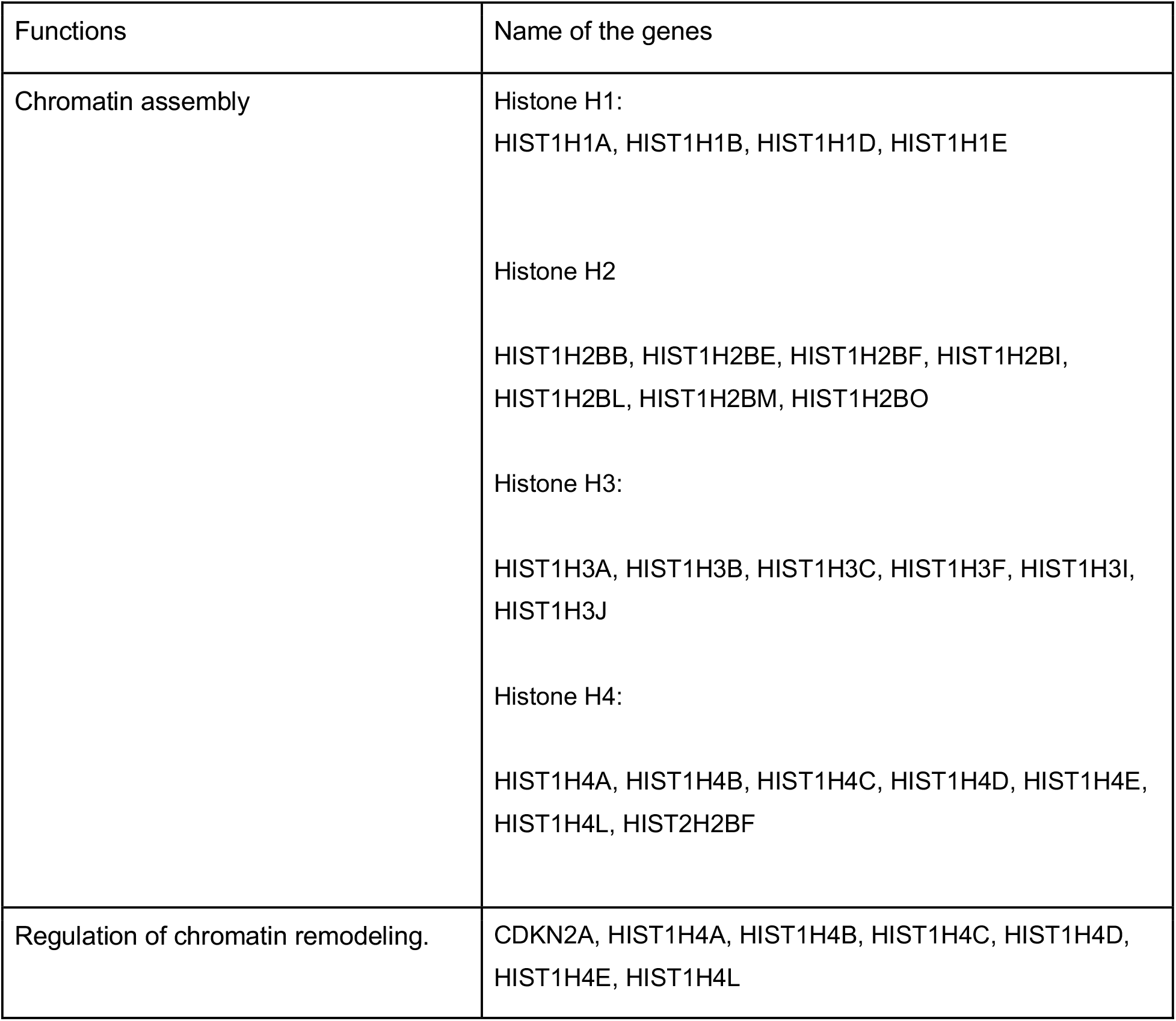
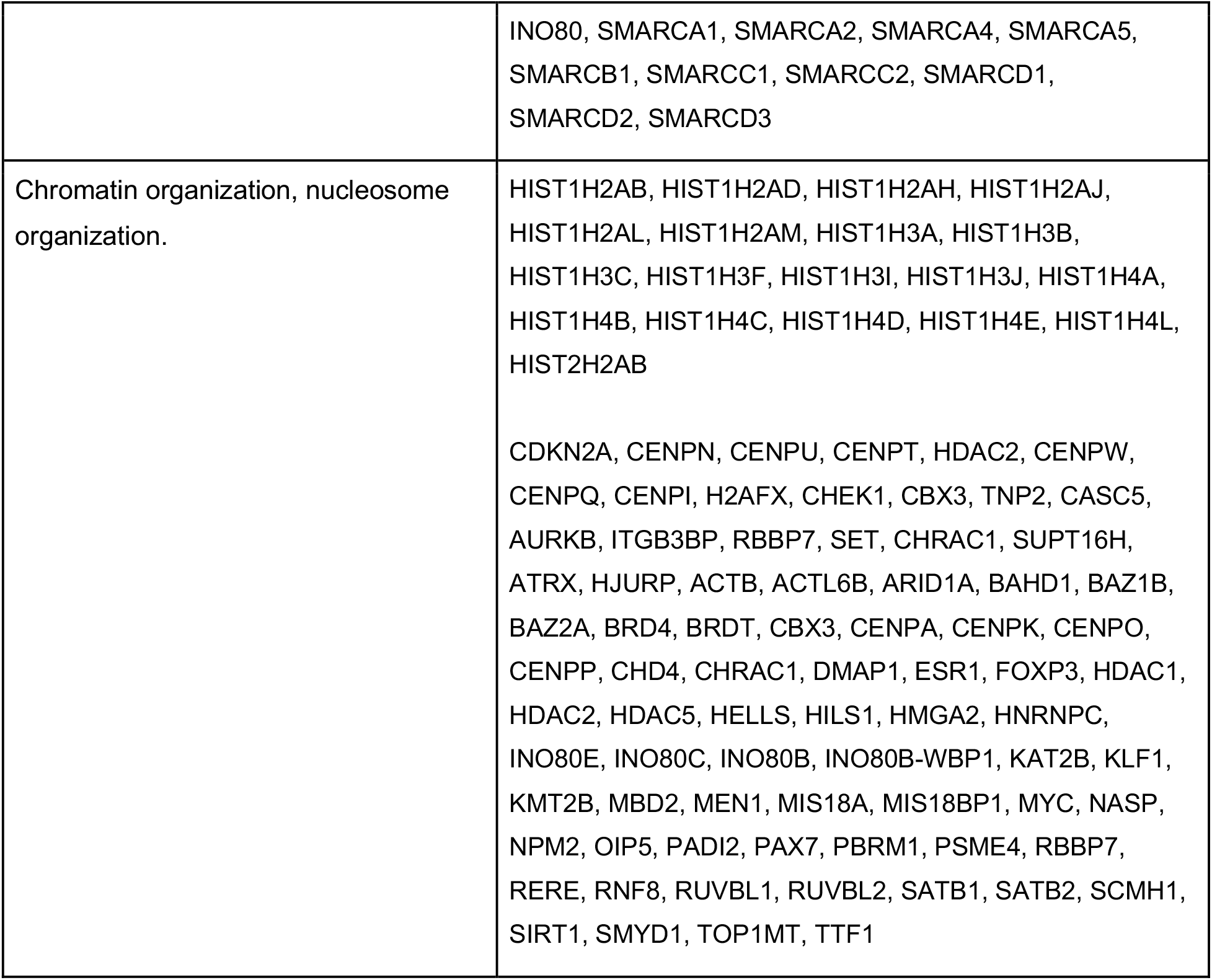
List of genes associated with chromosome organizations and assembly.

**Figure 5.**
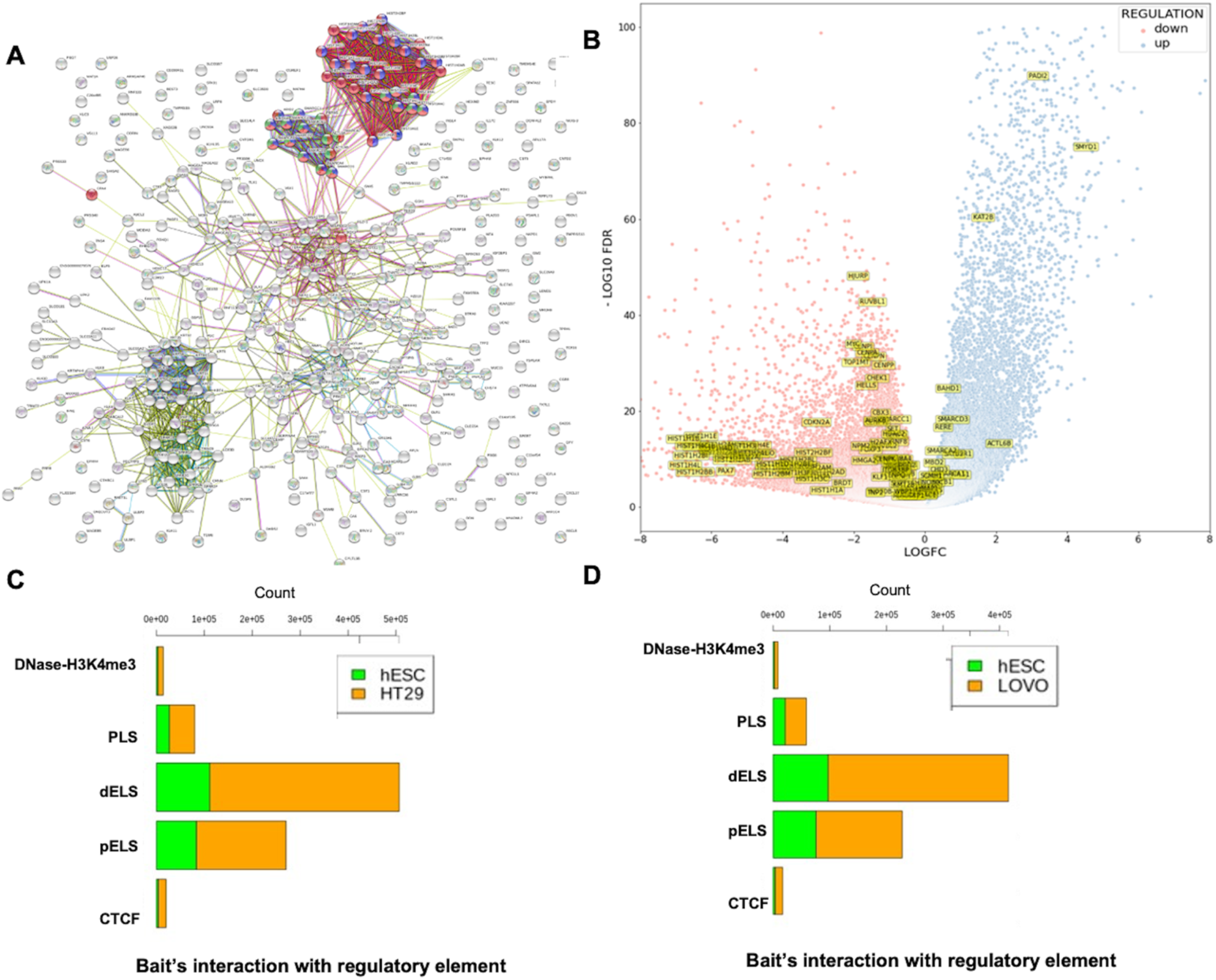
Functional annotation of genes extracted from TAD disrupted regions and proportion of cis-regulatory interaction with bait fragments. (A) Network visualization of genes and gene family associated with chromatin remodeling^95^ (Green), chromatin assembly (Blue), and chromatin organization (Red). (B) Volcano plot showing the expression pattern of chromatin remodeling, chromatin assembly and chromatin organization associated genes which had been extracted from gene ontology analysis. (C) The total number of bait interaction with regulatory elements (Promoter-like (PLS), Proximal enhancer-like (pELS), Distal enhancer-like (dELS), DNase-H3K4me3 and CTCF regions) in disrupted TADs regions of hESC and HT29 cell lines in hESC/HT29. (D) The total number of bait interaction with regulatory elements (Promoter-like (PLS), Proximal enhancer-like (pELS), Distal enhancer-like (dELS), DNase-H3K4me3 and CTCF regions) in disrupted TADs regions of hESC and LoVo cell lines in hESC/LoVo.

Further, 585 genes exhibited mutations specifically in LoVo and HT29 cells in which nonsynonymous and missense mutations were prominent. The frequency of mutations in affected genes involved in chromatin remodeling is depicted in Supplementary figure S2. The mutations along with amino acid changes are provided in the supporting information. Supplementary figure S2(A-B) provides the frequency of mutations in terms of CDs and nucleotide changes of genes associated with chromatin-associated genes collected from gene ontology analysis which lies in TAD disrupted regions based on Tate et al. and Chisanga et al.^73,74^. Supplementary figure S2(C-E) lists the various types of mutations in genes collected from the database^74^ which lie in the TAD disrupted regions in our case versus control study. We performed a comparative analysis with the list of chromatin remodeling associated genes collected from the gene ontology analysis coming from the TAD disrupted regions with relevant literature related to CRC^74–76^. We obtained 33, 41, and 106 common genes while comparing with the previous CRC studies^74–76^, respectively (Supplementary figure S3). Finally, we obtained 10 unique genes which were commonly affected in all CRC studies^74–76^ as well as in our study. Those genes are TOP1MT, HELLS, AURKB, CHEK1, SMARCC1, TTF1, HDAC1, SCMH1, RERE and PADI2, out of which HDAC1, RERE and PADI2 are known oncogenes^77^ (Supplementary figure S3).

### Distribution of the captured promoters’ interactions with candidate cis-regulatory elements (cCREs) derived from ENCODE data

The activation associated chromatin modification mark H3K4me3 makes the DNA in the chromatin more accessible for transcription factors, allowing the genes to be transcribed and expressed in the cell^78^. Similarly, distal enhancers activate target genes through DNA looping, a mechanism that enables distally bound transcription factors to contact the transcription machinery of target promoters^79–81^. As discussed in an earlier section, we used ChiCAGO, a PCHi-C interaction tool for finding bait interaction with other end regions using PCHi-C data of control (hESC) sample and case (HT29 and LoVo) samples (Figure 2). Each bait fragment was assigned a unique id which is termed as ‘bait id’ and the total number of interactions of each bait fragment with other ends (OE) regions termed as ‘bait count’. We only selected those bait ids having a non-zero bait count and bait count differences greater or equal than ten in the control versus case condition. Following the above criteria, we obtained 115198 and 630095 bait interactions with other ends (OE) regions of hESC and HT29 in the hESC/HT29 (Supplementary figure S4A) sample. Similarly, we obtained 112383 and 554536 bait interactions with other ends (OE) regions of hESC and LoVo cells respectively in the hESC/LoVo (Supplementary figure S4B) sample. For the sake of convenience, we referred to the above process as STAGE 1. Again, we filtered the bait ids from STAGE1 that have common bait ids generated in the previous section (Figure 3A). We referred to this filtered interaction as STAGE 2. Finally, we filtered bait ids of STAGE 2 that has common bait ids with the bait ids of the affected genes due to TAD disruption (Figure 4B). The bait ids of the list of affected genes can be filtered from baitmap IDs. The predicted digestion pattern of genomic DNA using the restriction enzyme site HindIII throughout the human genome (hg38) was generated using HiCUP tools. The final bait interactions file is referred as STAGE 3. We overlapped the other ends (OE) region of STAGE 3 interaction files with regulatory elements such as promoter-like (PLS), proximal enhancer-like (pELS), distal enhancer-like (dELS), DNase-H3K4me3 and CTCF regions (https://screen-v2.wenglab.org)^78^. It has clearly seen that there is high proximal enhancer-like (pELS) and distal enhancer-like (dELS) interaction with promoter capture counts compared to the remaining cis regulatory interactions (Figures 5C-D). However, the stagewise details of the total number of bait interactions with regulatory elements of hESC/HT29 and hESC/LoVo are given in Supplementary table S5.

### Role of enhancer-promoter interaction within TAD disruption region identified disease-relevance target genes in CRC

The integration of PCHi-C with Hi-C is to explore the genomic landscape from loop level to TADs-level then integration with differentially expressed genes coming from TCGA database and from HT29 (case) versus NT2D1 (control) cell lines to understand gene regulation. We overlapped with histone modification marks (H3K4me1 and H3K27ac) to verify the promoter interaction with enhancer which overall mimics the impact of TAD disruption on a gene’s expression. These could serve as key genes for potential therapeutic targets. Based on the capture count and log2fold value, we selected a few statistically significant DE genes that lie in the TAD disruption regions. The statistically significant gene list includes long non-coding (MALAT1, NEAT1, FTX and PVT1), small nucleolar (SNORA26 and SNORA71A) and protein-coding (TMPRSS11D, TSPEAR and DSG4). In this study, statistically significant implies we took only the top hit genes based on high capture counts, having enhancer-promoter loops, lie in TADs disruption regions and upregulated genes in CRC. For the visualization of Hi-C and PCHi-C interaction maps in the context of gene regulation, we highlighted one of the statistically significant upregulated gene locus *DSG4* in Figure 6A-B and remaining loci in Supplementary figure S5-11. The *DSG4* gene lies in the TAD disruption region in case (HT29 and LoVo) versus control cells (hESC) identified by Hi-C analysis. Similarly, PCHi-C identified more interaction frequencies between the *DSG4* promoter and, H3K27ac and H3K4me1 histone modification marks in case (HT29 and LoVo) compared to the control cells (hESC). TADs-based analysis helps in defining a gene’s cis -regulatory landscape while high-resolution promoter interaction data provides the required resolution to precisely map enhancer-promoter interactions. There were more active enhancer marks and enhancer-promoter loop formation in case while comparing with control cells around *DSG4* gene locus, suggesting its upregulation (Figure 6A-B). PCHi-C in different cell types identified thousands of enhancer-promoter contacts and revealed extensive differences in promoter architecture between cell types and throughout differentiation^43–48,50^. The variation in enhancer-promoter interactions within TAD disruption regions (Figure 6A-B) suggests that disease-relevant cell types (HT29 and LoVo) are critical for successful interrogation of the gene regulatory mechanisms of disease loci.

**Figure 6.**
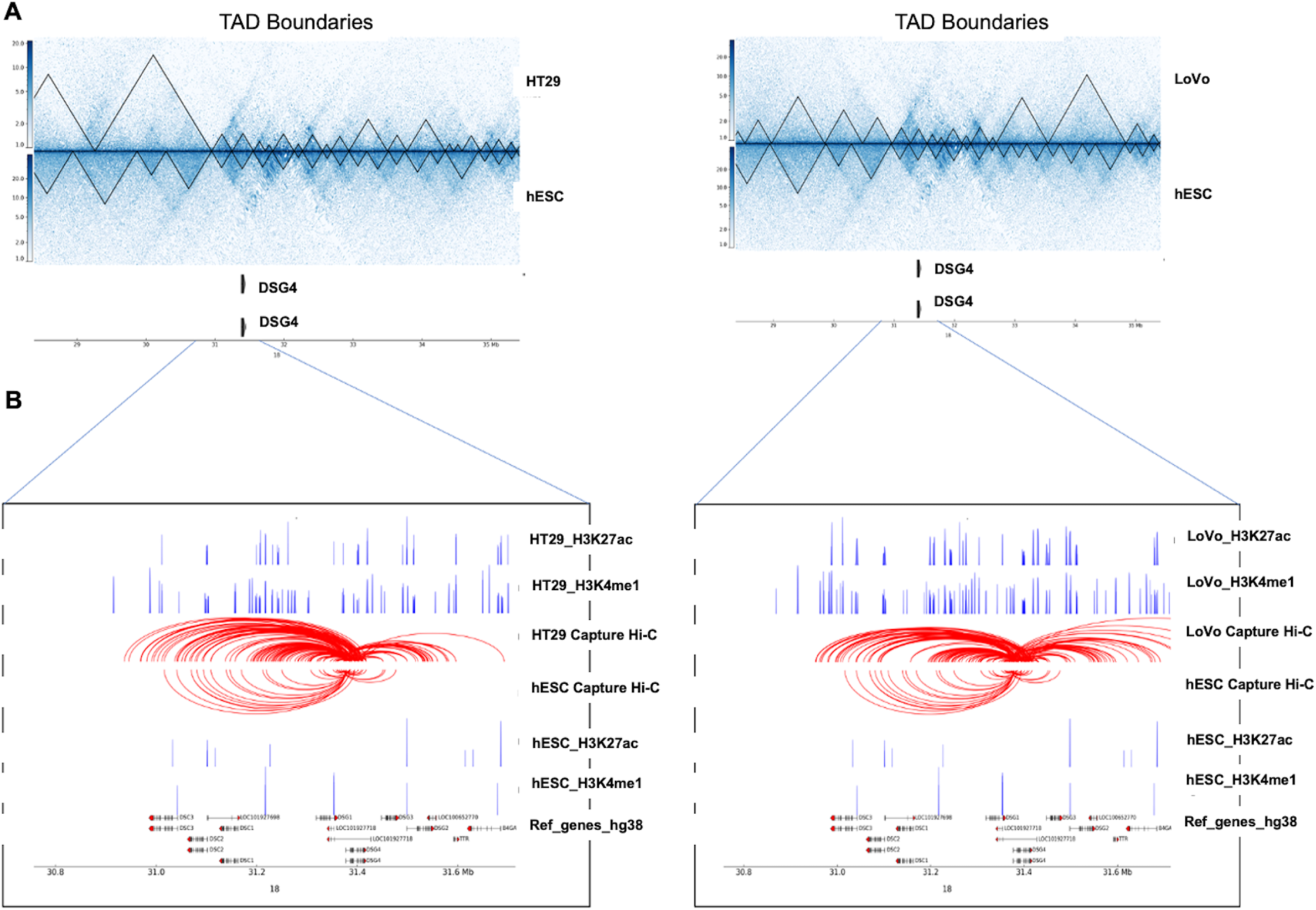
Effect on gene regulation due to structural changes between cancer (HT29 and LoVo) versus normal (hESC) cell lines. (A) ∼ 7 Mb region of chromosome 18 encompassing the *DSG4* gene is shown along with TAD boundaries of Hi-C interaction maps at 10 Kb resolution for case (HT29 and LoVo) and control (hESC). (B) Zoomed-in view of the *DSG4* locus in case (HT29 and LoVo) and control (hESC) along with corresponding PCHi-C interaction, and ChIP-seq data for H3K27ac, H3K4me1 are displayed in blue peaks. Filtered DSG4 read counts used by CHiCAGO are displayed in red with the corresponding significant interactions shown as arcs. For clarity, only DSG4 interactions were shown.

### Validation of the statistically significant genes revealed putative biomarkers

As discussed above, using our combined Hi-C, promoter-capture Hi-C, RNA-seq and ChIP-seq analysis, we have uncovered new potential targets which exhibit a high degree of dysregulation in CRC datasets with respect to the 3D structure. To validate whether the predictions of our analysis hold true, we next sought to monitor the expression profiles of a few of the statistically significant hits from our dataset in CRC models, some of which were previously reported to be dysregulated in CRC. lncRNA PVT1 has been implicated in the progression of colorectal cancer via the VEGFA-AKT axis^82^. lncRNA NEAT1 modulates chromatin accessibility in CRC and directly regulate Myc and ALDH1^83^. Similarly, recent studies^84–87^ have reported that the lncRNA FTX plays a crucial role in the initiation and progression of CRC. We used a highly malignant colorectal cell line HT29 and compared it to an embryonic cell line NT2D1 in our experimental validations to closely represent the primary datasets used for the bioinformatics analysis. We selected our targets in which the contact frequency was at least 2-fold. We observed that expression levels of genes belonging to long non-coding (MALAT1, NEAT1, FTX and PVT1), small nucleolar (SNORA26 and SNORA71A) as well as protein-coding families (TMPRSS11D, TSPEAR and DSG4) were significantly much higher in HT29 as well as various other colorectal model cell lines when compared to NT2D1 embryonic cells (Figure 7A). Next, we monitored the relative levels of transcriptional activation associated chromatin modifications H3K4me3 and H3K27ac at the promoters of these genes by performing ChIP-quantitative PCRs. We observed a significant enrichment of both these modifications at the promoters of all selected target genes (Figure 7B-C), suggesting a hierarchical modification of the chromatin following TAD disruption. Collectively, these analyses suggest that changes in transcription associated chromatin modifications correlated with the 3D chromatin changes shown previously, hence leading to activation of the transcription machinery.

**Figure 7.**
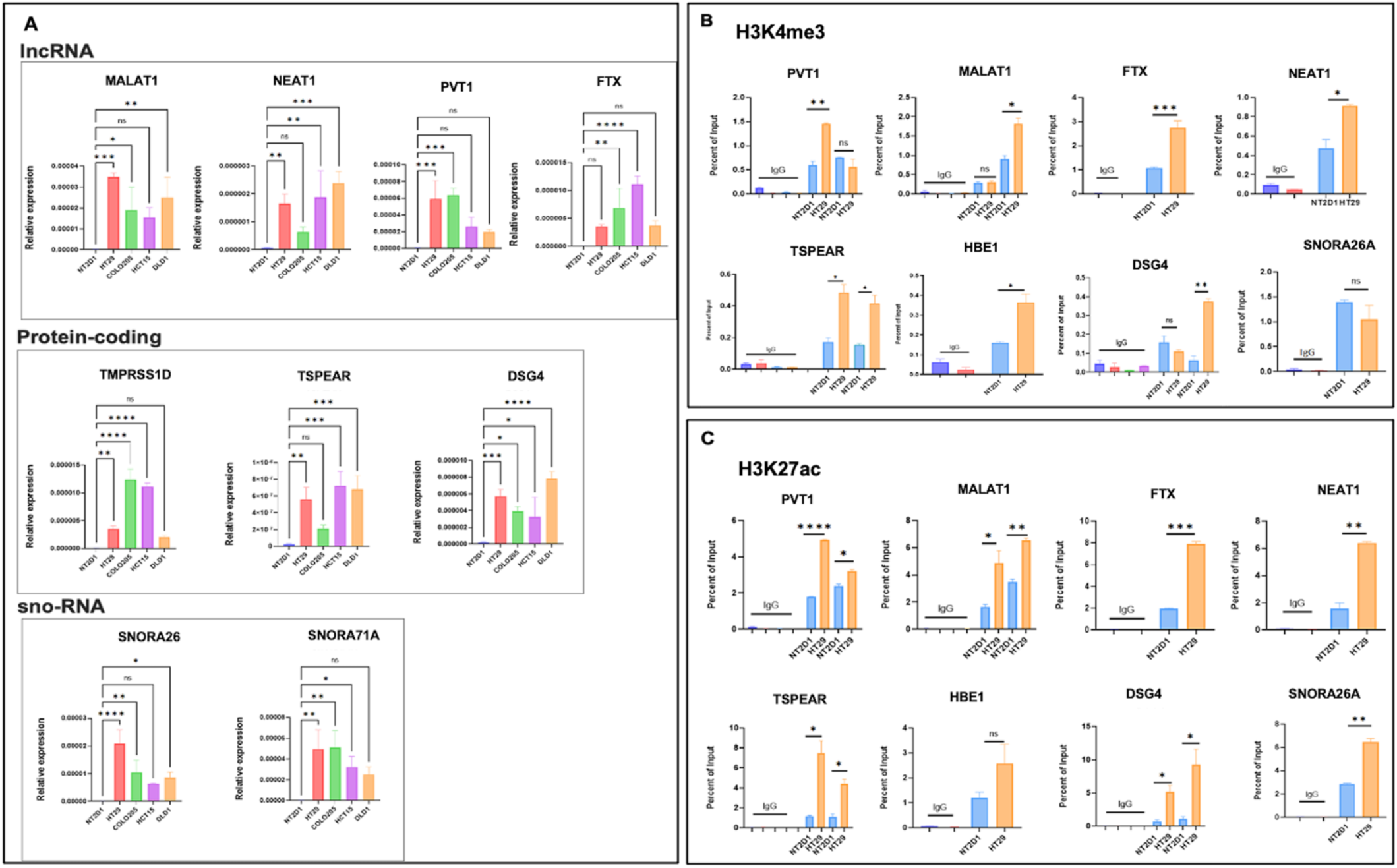
Validation of target genes extracted from integrative analysis. (A) qRT-PCR analysis of target genes in NT2D1, HT29, Colo205, HCT15 and DLD1 cell lines, showing relative enrichment of gene expression in the CRC cell lines compared to NT2D1. (B and C) ChIP-qPCR for H3K4me3 and H3K27ac modifications, respectively, on the transcription start site (TSS) upstream regions of target genes. Three biological replicates were used for statistical analysis using Graphpad v9.1. One-way ANOVA or unpaired t-test was used as the significance test. * p<0.05, ** p<0.01, *** p<0.001, **** p<0.0001, ns-non-significant.

### Expression pattern of statistically significant genes for therapeutic target in single cell human colon cancer atlas

The integrative analysis reported above is based on bulk-seq data. In bulk-seq study, samples are treated like homogeneous population. However, the cell populations in the human body are heterogeneous in nature and each cell reflects unique activity. Due to recent breakthroughs, it is now possible to analyze the transcriptome at single-cell level for over millions of cells^88,89^. This enabled us to differentiate, characterize and classify each cell at the transcriptome level, which leads to prediction of rare cell population/s. Taking the advantage of the single cell human colon cancer atlas database^90^, we analyzed transcriptionally profiled 371,223 tumor and adjacent normal cells generated based on scRNA-seq. In the analysis step, first we preprocessed data for preventing outlier cells which could influence downstream analysis. We generated a violin plot before and after performing the quality control (Figure 8A-B). In the quality control step, we manually selected the threshold level of number of features (transcripts), number of counts against each feature and mitochondrial percentage of cells. Using the processed data, we generated a t-SNE plot (Figure 8C). The t-SNE plot separated the diverse cell population into 36 clusters enabling high-resolution depiction of the cellular diversity and heterogeneity. Here, we were interested in monitoring the expression patterns derived from the integrative bulk-seq study at single cell level across different cell populations. This helped us to gauge the distribution of the expression patterns of targeted genes across diverse cell populations (Figure 8C). In this study we selected a few statistically significant genes (MALAT1, NEAT1, FTX, PVT1, SNORA26, SNORA71A, TMPRSS11D, TSPEAR and DSG4) as potential therapeutic targets in early colorectal cancer detection or prevention. Out of these 9 genes, 3 genes (SNORA26, SNORA71A, TMPRSS11D) were missing in the single cell gene annotation files. Therefore, we monitored the expression pattern of the remaining six genes using the single cell human colon cancer atlas database^90^. FTX and PVT1 exhibit lower expression levels compared to MALAT1 and NEAT1 (Figure 8D). FTX gene showed significant expression only in B cell, T cell, endothelial cell, epithelial cell, macrophage, and monocyte populations. PVT1 gene showed significant expression only in B cell, T cell, and epithelial cell cells population. However, TSPEAR and DSG4 did not exhibit any significant expression level differences among distinct cell populations. Expression profiling at single cell level helped us to monitor the distribution of gene expression across distinct cell types which could be considered as a key factor in the assessment of their potential as a biomarker.

**Figure 8.**
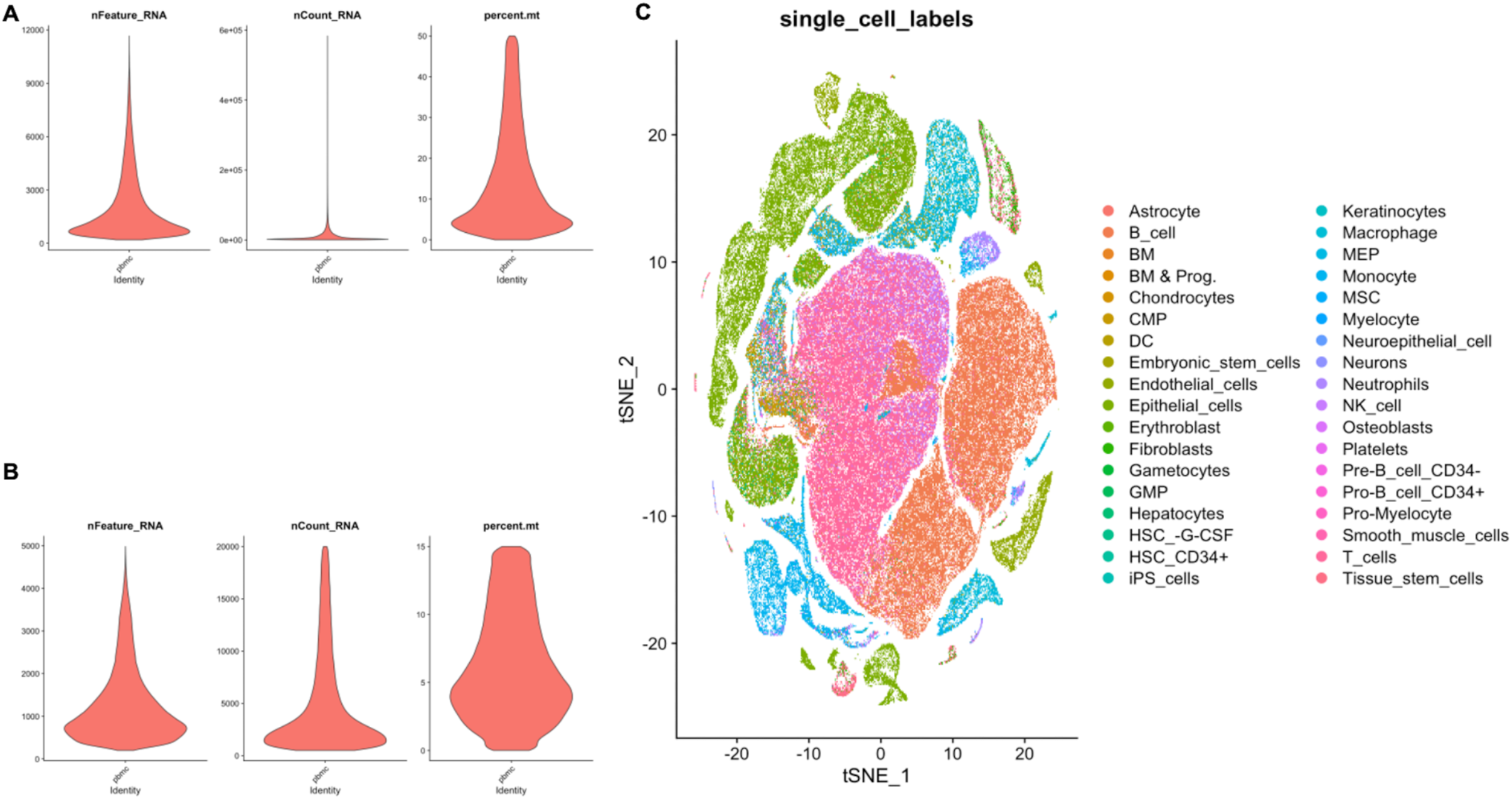
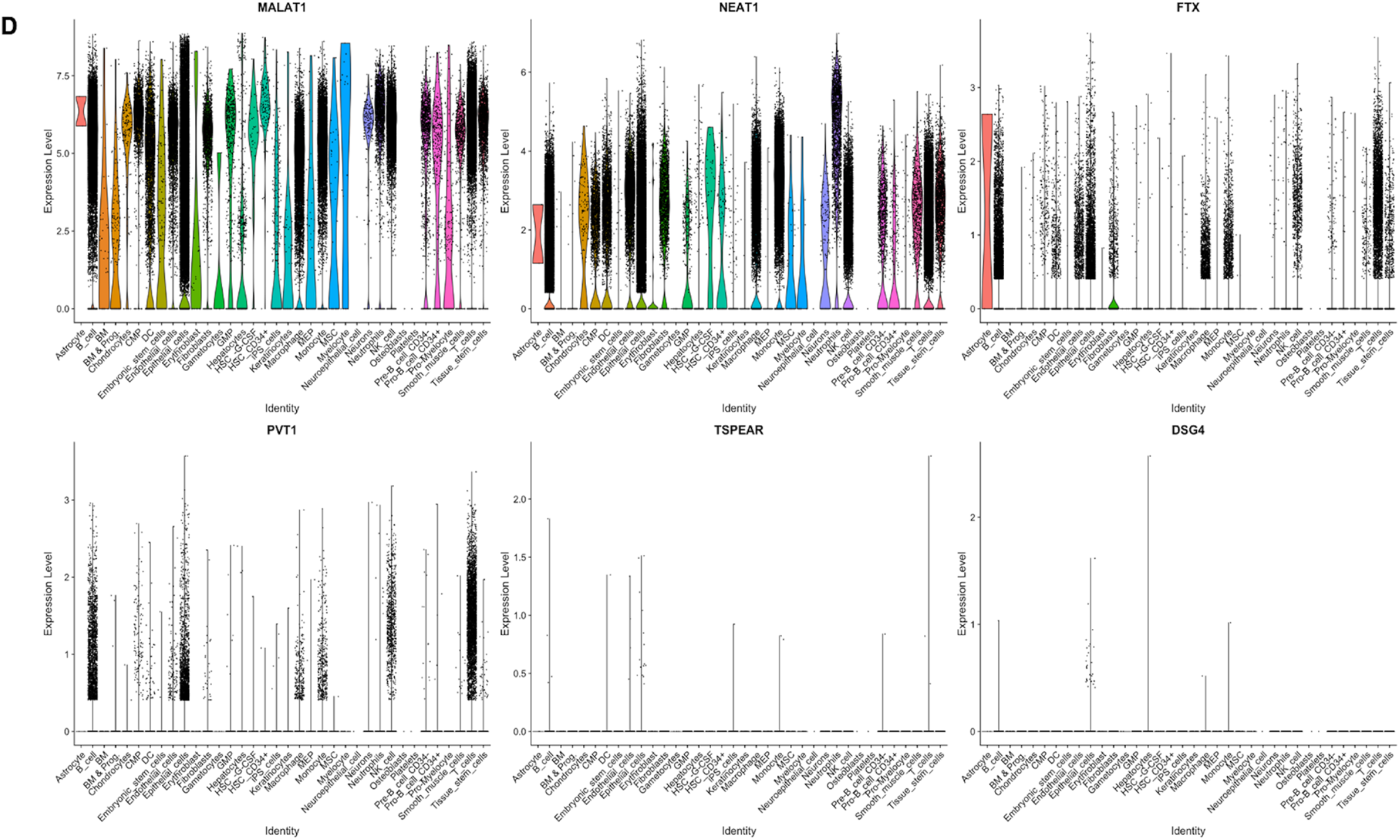
Single-cell RNA-Seq analysis of 371,223 tumor and adjacent normal cells collected from single cell human colon cancer atlas database reveals contributions of target genes across diverse cell populations. (A and B) Violin plot of number of features, number of counts and mitochondrial percentage before and after quality control. (C) t-SNE plot of 371,223 tumor and adjacent normal cells separated the cell population into 36 clusters enabling high-resolution depiction of the cellular diversity and heterogeneity. (D) Violin plots of targeted gene enrichment scores across 36 diverse cell type populations.

## DISCUSSION

TADs are emerging as the major feature of 3D chromosome organization and gene regulation. We aimed to dissect how these domains are formed and regulate enhancer-promoter communication and gene expression. In this study we explored how 3D folding defects can lead to altered gene expression in colorectal cancer. Due to genome instability in cancer genomes, TAD disruption occurs more frequently in cancer compared to normal cells which can lead major changes in gene expression that may possibly drive tumorigenesis. Our comprehensive analysis attempted to explore the molecular basis of the genetic risk for CRC. The patterns of TADs in cancer cells may therefore contribute to a better understanding of the genetic basis of cancer cell gene expression, and possibly provide specific targets for treatment. Therefore, identifying multiple target genes which may potentially serve as therapeutic targets for colorectal cancer, we presented a systematic integration of genome topology, loop interaction and gene regulation in colorectal cancer using publicly available Hi-C, PCHi-C, ChIP-seq, RNA-seq and scRNA-seq datasets. Generating high-resolution Hi-C data by conducting Hi-C experiments needs millions of mammalian cells, which may generate billions of paired-end reads. Due to high sequencing cost, the resolution of most Hi-C datasets is coarse, and it is not ideal for looking at fine-scale interactions around the regions of interest such as disease-associated loci. To overcome this limitation, we integrated PCHi-C data with Hi-C to understand the dynamics of regulatory chromosomal interactions from local-to-TAD level. During such analysis, we noticed most interactions were found in cancer cell lines (HT29, LoVo) compared to hESCs indicating higher proliferation in tumor cell population compared to human embryonic stem cells. TADs are conserved during evolution and play critical roles in controlling and driving long-range regulation of gene expression^91^. TADs disrupted by deleting a TAD boundary leads to fusion of the two adjacent TADs or genomic rearrangements that break up the TADs and creates new TADs without affecting the TAD boundaries. Disruption of TAD boundaries results in aberrant gene expression by exposing genes to inappropriate regulatory elements. TAD disruption is often found in cancer cells and contributes to oncogenesis^92–94^. Understanding the role of TAD disruption and long-range chromatin interactions will help us to provide insights into the mechanism of gene regulation in general and will also reveal how genomic rearrangements and mutations in cancer genomes can lead to the aberrant expression of oncogenes and tumor suppressors. For gene expression analysis, we performed differential RNA-seq analysis using colorectal cancer samples (primary tumor, metastatic) compared with normal samples (blood-derived normal, solid tissue) obtained from the TCGA databases. We filtered differentially expressed genes found around variably interacting regions to check the distribution of DGEs across TADs boundaries. We used H3K27ac and H3K4me1 histone modification marks for understanding the gene’s cis-regulatory landscape to precisely map the enhancer-promoter interactions. The functional study revealed that most of the genes upregulated in cancer were found around TAD disruption regions, and are involved in chromatin remodeling, chromatin assembly and chromatin organization. We also integrated the mutational datasets collected from databases like cosmic database^73,74^ and human oncogene database^77^ with the affected genes due to TAD disruptions. Results of such analyses showed that key genes AR1D1A, ATRX, centromere protein complex, histone complex, SATB1, SATB2, SWI1/SNF1 complex, histone deacetylases were frequently mutated in these datasets. Finally for experimental validation, we filtered statistically significant gene-list based on high capture count and log_2_fold change in transcript expression; and with relevant literature related to colorectal cancer. We found expression of long non-coding genes (MALAT1, NEAT1, FTX and PVT1), small nucleolar (SNORA26 and SNORA71A) and protein-coding (TMPRSS11D, TSPEAR and DSG4) were significantly much higher in HT29 compared to NT2D1; and enriched with activation-associated histone modifications H3K4me3 and H3K27ac. Our data, while confirming the expression patterns, strongly argues in favor of *a priori* involvement of epigenetic mechanisms in form of dynamic histone modifications as well as the chromatin interaction changes that modulate the gene expression. Additionally, for understanding the expression patterns of target genes across diverse cell types, we analyzed 371,223 tumor and adjacent normal cells taken from single cell human colon cancer atlas database. We found expression of FTX and PVT1 is contributed from the rare cell population while MALAT1 and NEAT1 were expressed almost everywhere in diverse cell populations. However, expression of TSPEAR and DSG4 was not observed in the scRNA-seq analysis. Collectively, our study suggests that changes in 3-D genomic architecture affect transcriptome signatures which might be associated with tumor-suppressive transcriptional programs. Based on this analysis we identified multiple target genes which may potentially serve as biomarkers and could be used to better understand CRC progression.

### Limitations of the study

Chromatin loops constitute an important feature in the structural organization of the genome. Due to lower sequencing depth of the available Hi-C datasets, we were not able to detect fine-scale interactions at genome-wide scale using the loop detection tool. Therefore, to overcome these issues, we used promoter capture Hi-C (PCHi-C) data for mapping the fine-scale interactions. In Capture Hi-C data, the captured regions were selected based on genome-wide association studies (GWAS) which contains of 34,627 CRC cases and 71,379 controls of European ancestry that identifies SNPs at 31 new CRC risk loci. Therefore, this study is limited to 31 CRC risk loci instead of all genomic loci to monitor the genomic structural instability of colorectal cancer cells (HT29 and LoVo) compared to hESCs. Further validation using primary CRC tumor samples is required to ascertain the clinical significance of these findings.

## Supporting information

Saw et al., Supplementary_table_list

## DATA AVALIABILITY

Hi-C and PCHi-C sequencing data for HT29, LoVo and hESC cell lines have been downloaded from the European Genome-phenome Archive (EGA) under the accession number EGAS00001001946 and NCBI GEO under the accession number GSM892306 and GSM2309026. The H3K27ac and H3K4me1 ChIP-seq samples of HT29, LoVo and hESC cell lines were downloaded from NCBI GEO under the SRA number SRR5063155, SRR5063156, SRR5063159, SRR5063160, SRR2481801, SRR8541250 and European Genome-phenome Archive (EGA) under the accession code EGAS00001001946. HTSeq-Counts of gene expression data (45 normal samples and 568 tumor samples) were collected from The Cancer Genome Atlas (TCGA) database (https://www.cancer.gov/tcga). The RNA-seq samples of HT29 and NT2 cell lines were downloaded from NCBI GEO under the SRA number SRR592573, SRR592574, SRR592575, SRR2075688 and SRR2075689. The scRNA-seq of 371,223 tumor and adjacent normal samples were downloaded from NCBI GEO under accession number GSE178341. Additional resources related to scRNA-seq data are available in the web page (https://broad.io/crchubs) and the Broad Institute’s Single Cell Portal (https://singlecell.broadinstitute.org/single_cell/study/SCP1162).

## SUPPLEMENTARY DATA

Contains 12 supplementary figures and 5 tables.

## CODE AVALIABILITY

All bioinformatics and statistical analysis tools used are open source. Custom scripts of the generated results and figures in the manuscript are available upon request.

## ACKNOWLEDGEMENT

AKS acknowledges SERB for National Postdoctoral Fellowship. AM acknowledges fellowship from the Council of Scientific and Industrial Research (CSIR), Government of India. SN acknowledges European Genome-phenome Archive (EGA) for sharing the CRC data. The support and the resources provided by ‘PARAM Brahma Facility’ under the National Supercomputing Mission, Government of India at the Indian Institute of Science Education and Research (IISER) Pune are gratefully acknowledged. SG acknowledges Science and Engineering Research Board, Government of India for the JC Bose Fellowship (JCB/2019/000013).

