## Supplementary Figures 1-11 for "Combined promoter-capture Hi-C and Hi-C analysis reveals a fine-tuned regulation of 3D chromatin architecture in colorectal cancer"

[*] To whom correspondence should be addressed.

A

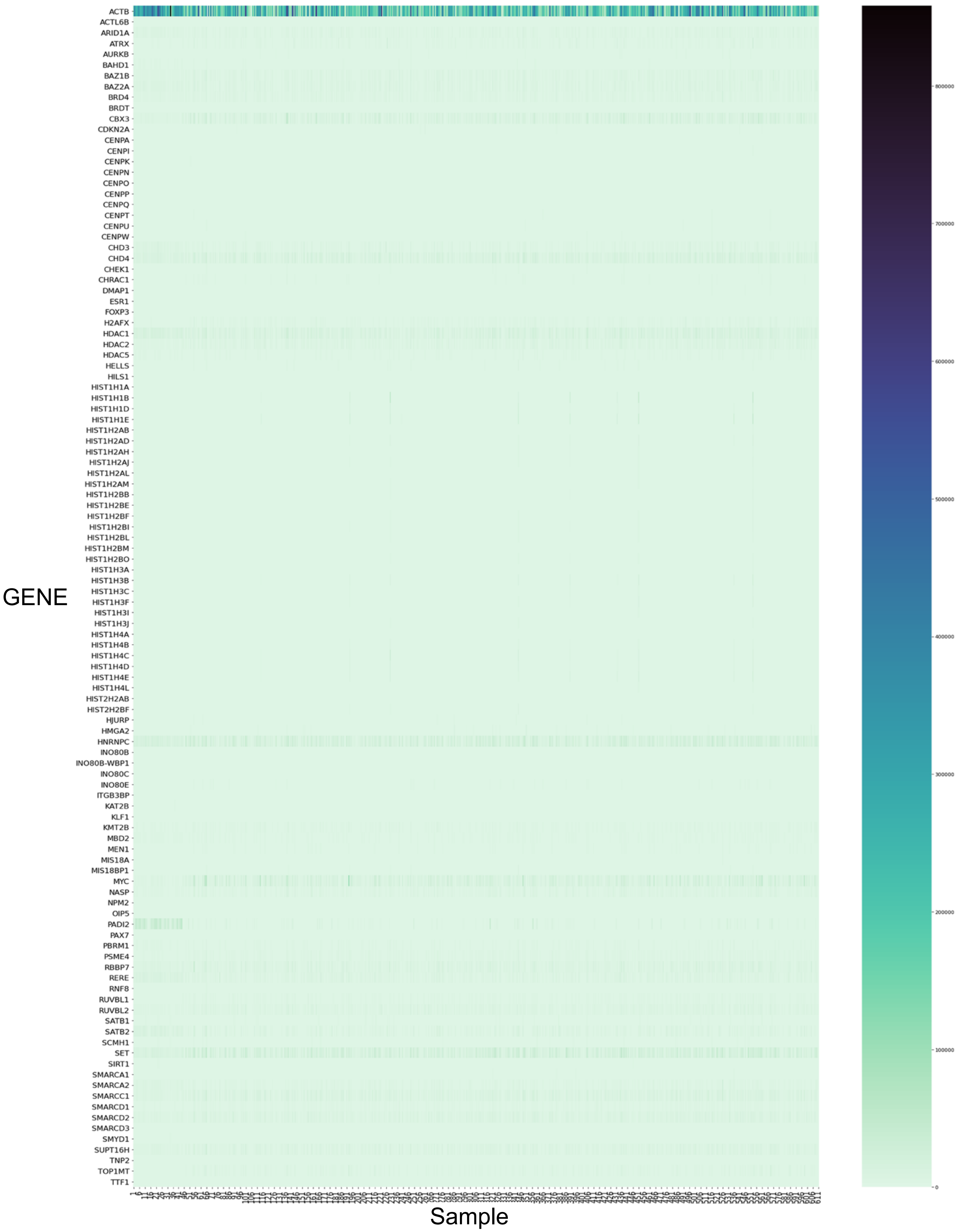
ACTB
ACTL6B
ARID1A
ATRX
AURKB
BAHD1
BAZ1B
BAZ2A
BRD4
BRDT
CBX3
CDKN2A
CENPA
CENPI
CENPK
CENPN
CENPO
CENPP
CENPQ
CENPT
CENPU
CENPW
CHD3
CHD4
CHEK1
CHRAC1
DMAP1
ESR1
FOXP3
H2AFX
HDAC1
HDAC2
HDAC5
HELLS
HILS1
HIST1H1A
HIST1H1B
HIST1H1D
HIST1H1E
HIST1H2AB
HIST1H2AD
HIST1H2AH
HIST1H2AJ
HIST1H2AL
HIST1H2AM
HIST1H2BB
HIST1H2BE
HIST1H2BF
HIST1H2BI
HIST1H2BL
HIST1H2BM
HIST1H2BO
HIST1H3A
HIST1H3B
HIST1H3C
HIST1H3F
HIST1H3I
HIST1H3J
HIST1H4A
HIST1H4B
HIST1H4C
HIST1H4D
HIST1H4E
HIST1H4L
HIST2H2AB
HIST2H2BF
HJURP
HMGA2
HNRNPC
INO80B
INO80B-WBP1
INO80C
INO80E
ITGB3BP
KAT2B
KLF1
KMT2B
MBD2
MEN1
MIS18A
MIS18BP1
MYC
NASP
NPM2
OIP5
PADI2
PAX7
PBRM1
PSME4
RBBP7
RERE
RNF8
RUVBL1
RUVBL2
SATB1
SATB2
SCMH1
SET
SIRT1
SMARCA1
SMARCA2
SMARCC1
SMARCD1
SMARCD2
SMARCD3
SMYD1
SUPT16H
TNP2
TOP1MT
TTF1
GENE
Sample
800000
700000
600000
500000
400000
300000
200000
100000
0

B

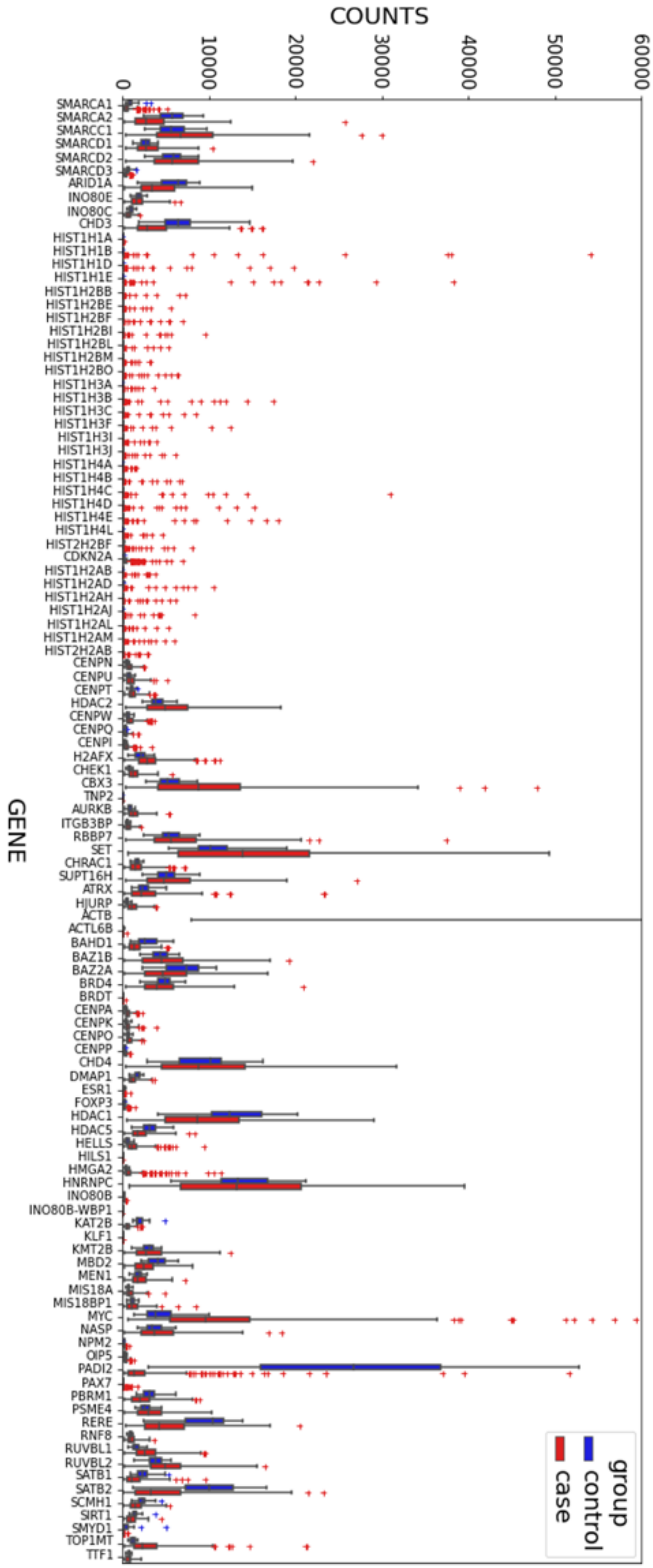


**Figure S1. GO ontology analysis.** (A) Heatmap and (B) Normalized count showing the expression pattern of chromatin remodeling, chromatin assembly and chromatin organization associated genes which has been extracted from gene ontology analysis.

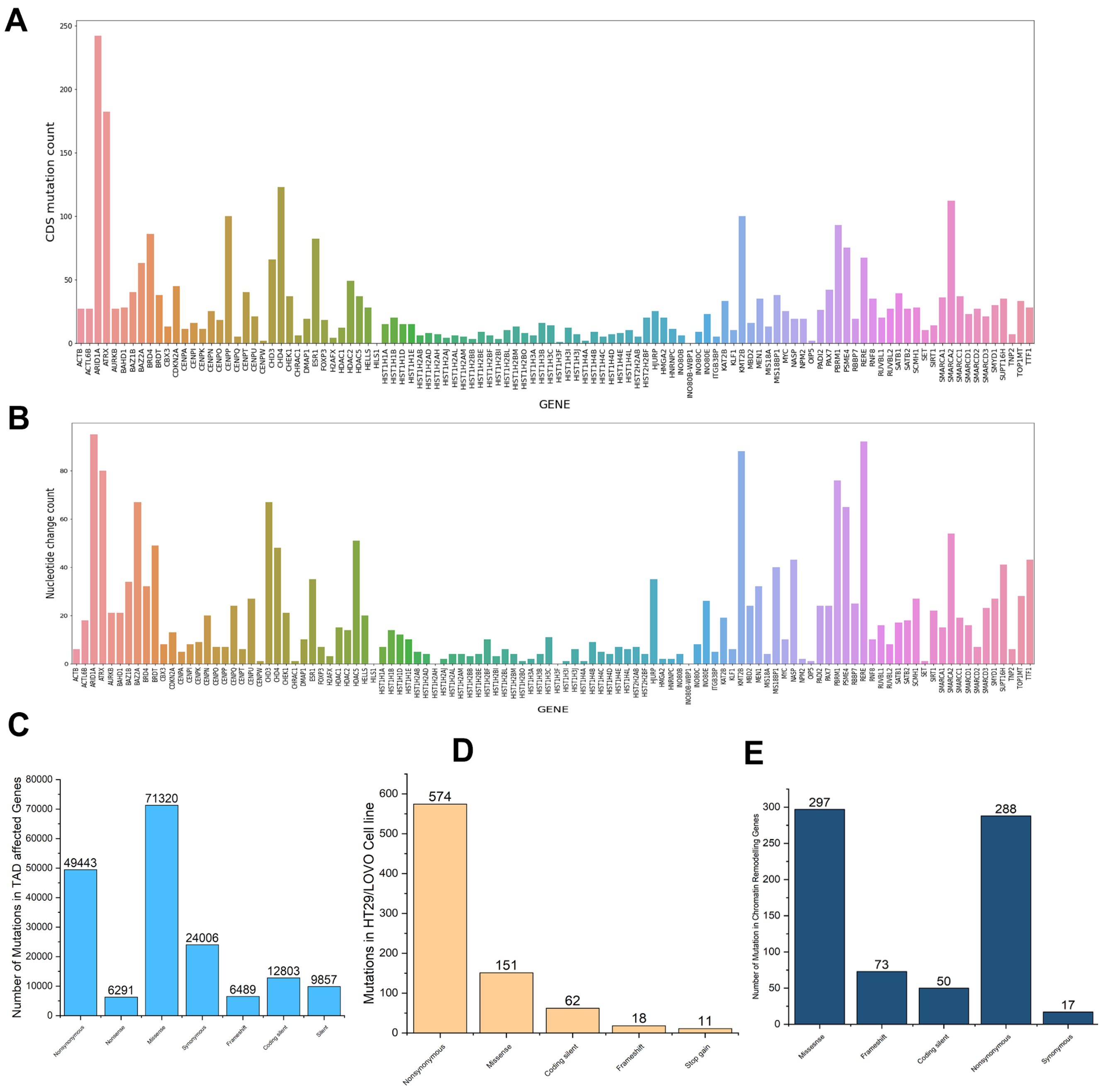


**Figure S2. The variation in number and type of mutation.** (A) Frequency of CDS mutations in genes associates with chromatin-associated genes collected from gene ontology analysis which lies in TAD disrupted regions[1]. (B) Frequency of mutations of nucleotide changes in chromatin-associated genes collected from gene ontology analysis which lies in TAD disrupted regions[2]. (C) Types of mutations in genes collected from colorectal cancer cell lines database[2] which lies in TAD disrupted regions in our case versus control study. (D) Types of mutations in genes from HT20 and LoVo cell lines[2] lies in TAD disrupted regions in our case versus control study. (E) Type of mutations in chromatin-associated genes collected from gene ontology analysis which lies in TAD disrupted regions[2] .

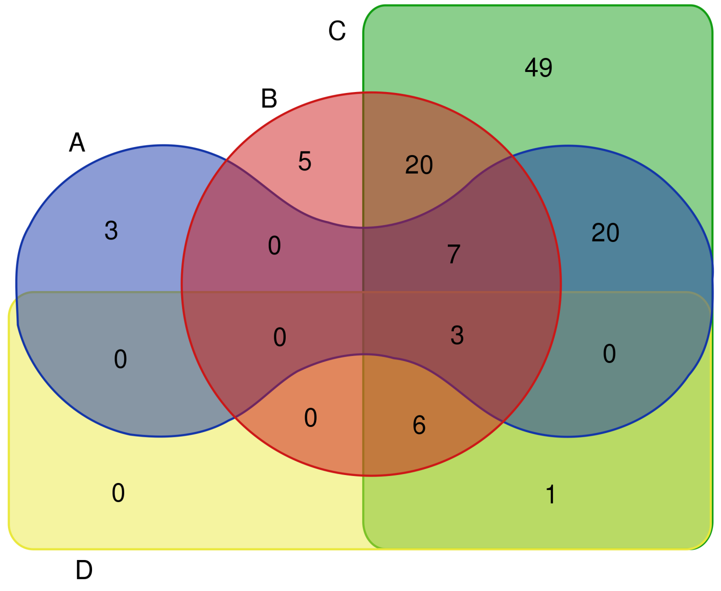


**Figure S3. Identification of potential genes in colorectal cancer susceptibility.** Venn diagram representing the list of genes in our study which also found in colorectal cancer relevant literature studies[2–5]. Here, number represent distinct gene count and four different colors corresponds to scientific articles[2-5] related to colorectal cancer study.

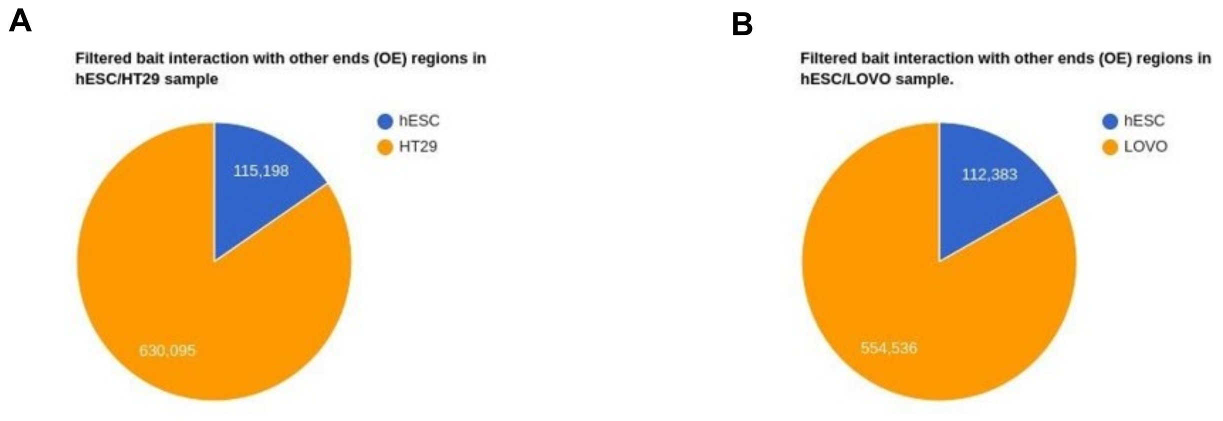


**Figure S4. The proportion of capture Hi-C interaction in normal versus cancer cell lines.** (A) Filtered bait interaction with other ends (OE) regions of hESC and HT29 in the hESC/HT29 sample. (B) Filtered bait interaction with other ends (OE) regions of hESC and LoVo in the hESC/LoVo sample.

### TAD Boundaries

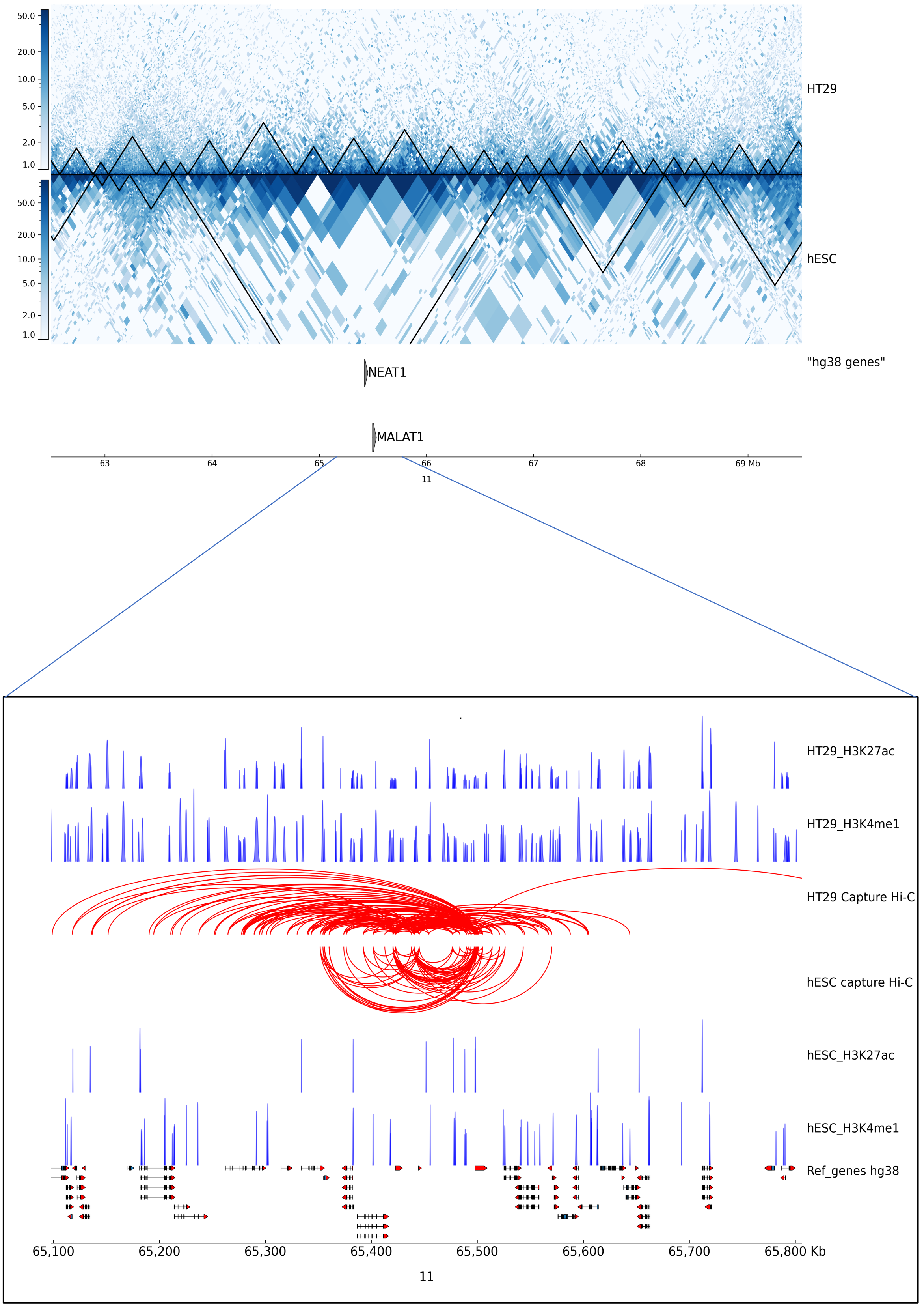

**Figure S5 (I). Effect on gene regulation due to structural changes between cancer (HT29) versus normal (hESC) cell lines. (**A) ~ 7 Mb region of chromosome 11 encompassing the *MALAT1* and *NEAT1* genes is shown along with TAD boundaries of Hi-C interaction maps at 10 Kb resolution for case (HT29) and control (hESC). (B) Zoomed-in view of the *MALAT1* and *NEAT1* loci in case (HT29) and control (hESC) along with corresponding PCHi-C interaction, and ChIP-seq data for H3K27ac, H3K4me1 are displayed in blue peaks. Filtered MALAT1 and NEAT1 read counts used by CHiCAGO are displayed in red with the corresponding significant interactions shown as arcs. For clarity, only *MALAT1* and *NEAT1* interactions are shown.

### TAD Boundaries

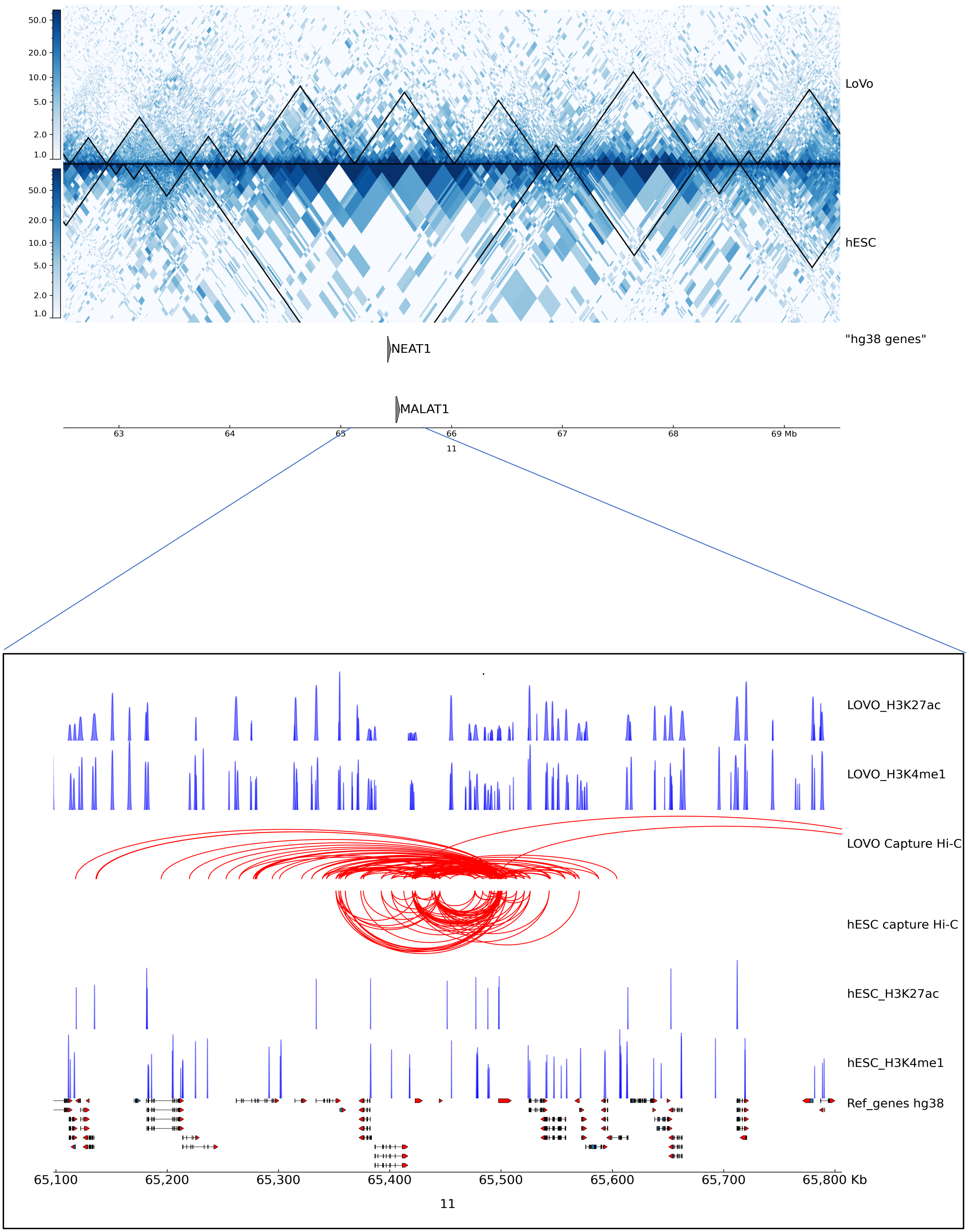

**Figure S5(II). Effect on gene regulation due to structural changes between cancer (LoVo) versus normal (hESC) cell lines.** (A) ~ 7 Mb region of chromosome 11 encompassing the *MALAT1* and *NEAT1* genes is shown along with TADs boundaries of Hi-C interaction maps at 10 Kb resolution for case (LoVo) and control (hESC). (B) Zoomed-in view of the *MALAT1* and *NEAT1* loci in case (LoVo) and control (hESC) along with corresponding PCHi-C interaction, and ChIP-seq data for H3K27ac, H3K4me1 are displayed in blue peaks. Filtered MALAT1 and NEAT1 read counts used by CHiCAGO are displayed in red with the corresponding significant interactions shown as arcs. For clarity, only *MALAT1* and *NEAT1* interactions are shown.

### TAD Boundaries

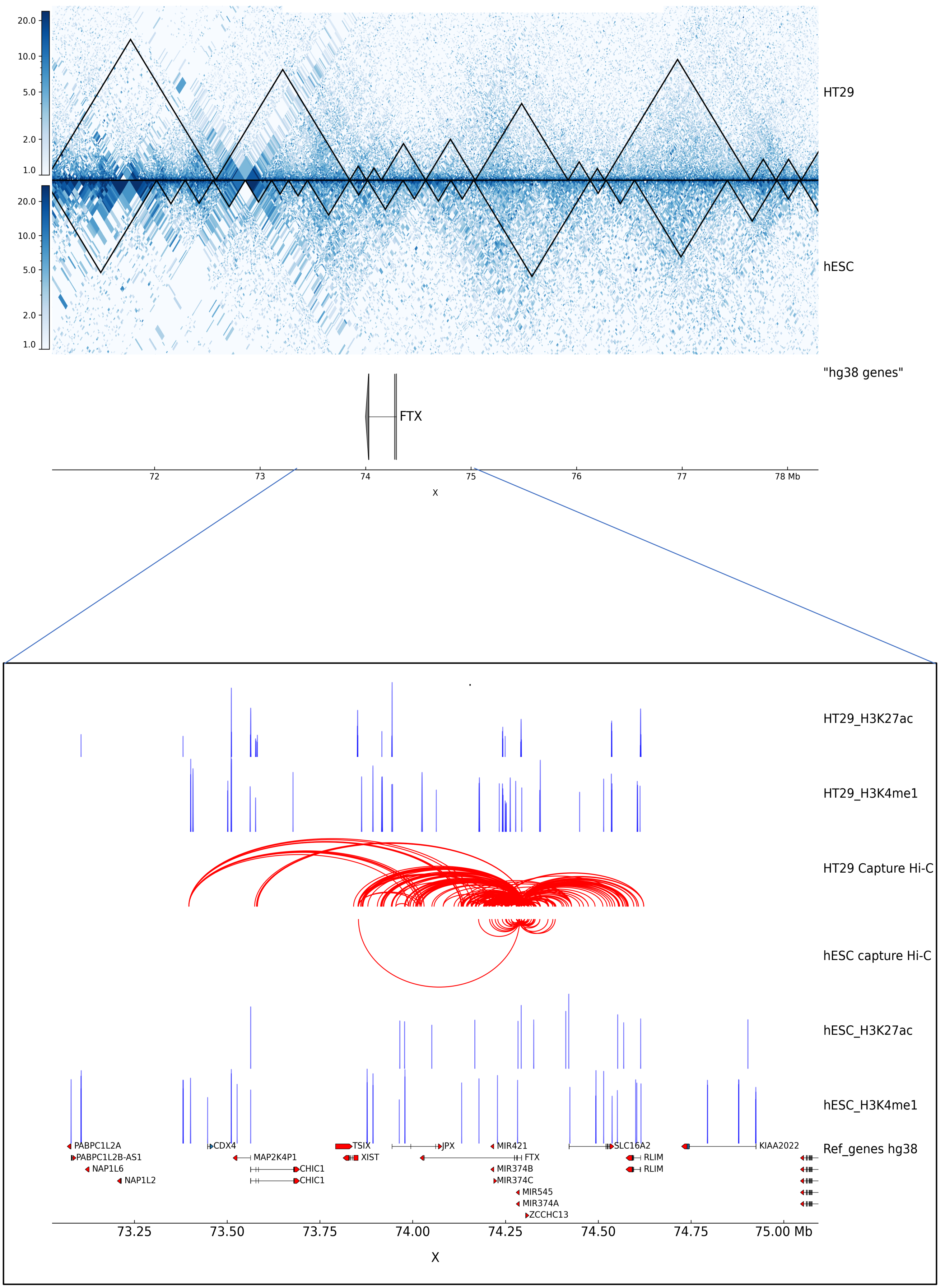

**Figure S6 (I). Effect on gene regulation due to structural changes between cancer (HT29) versus normal (hESC) cell lines.** (A) ~ 7 Mb region of chromosome X encompassing the *FTX* gene is shown along with TADs boundaries of Hi-C interaction maps at 10 Kb resolution for case (HT29) and control (hESC). (B) Zoomed-in view of the *FTX* locus in case (HT29) and control (hESC) along with corresponding PCHi-C interaction, and ChIP-seq data for H3K27ac, H3K4me1 are displayed in blue peaks. Filtered FTX read counts used by CHiCAGO are displayed in red with the corresponding significant interactions shown as arcs. For clarity, only *FTX* interactions are shown.

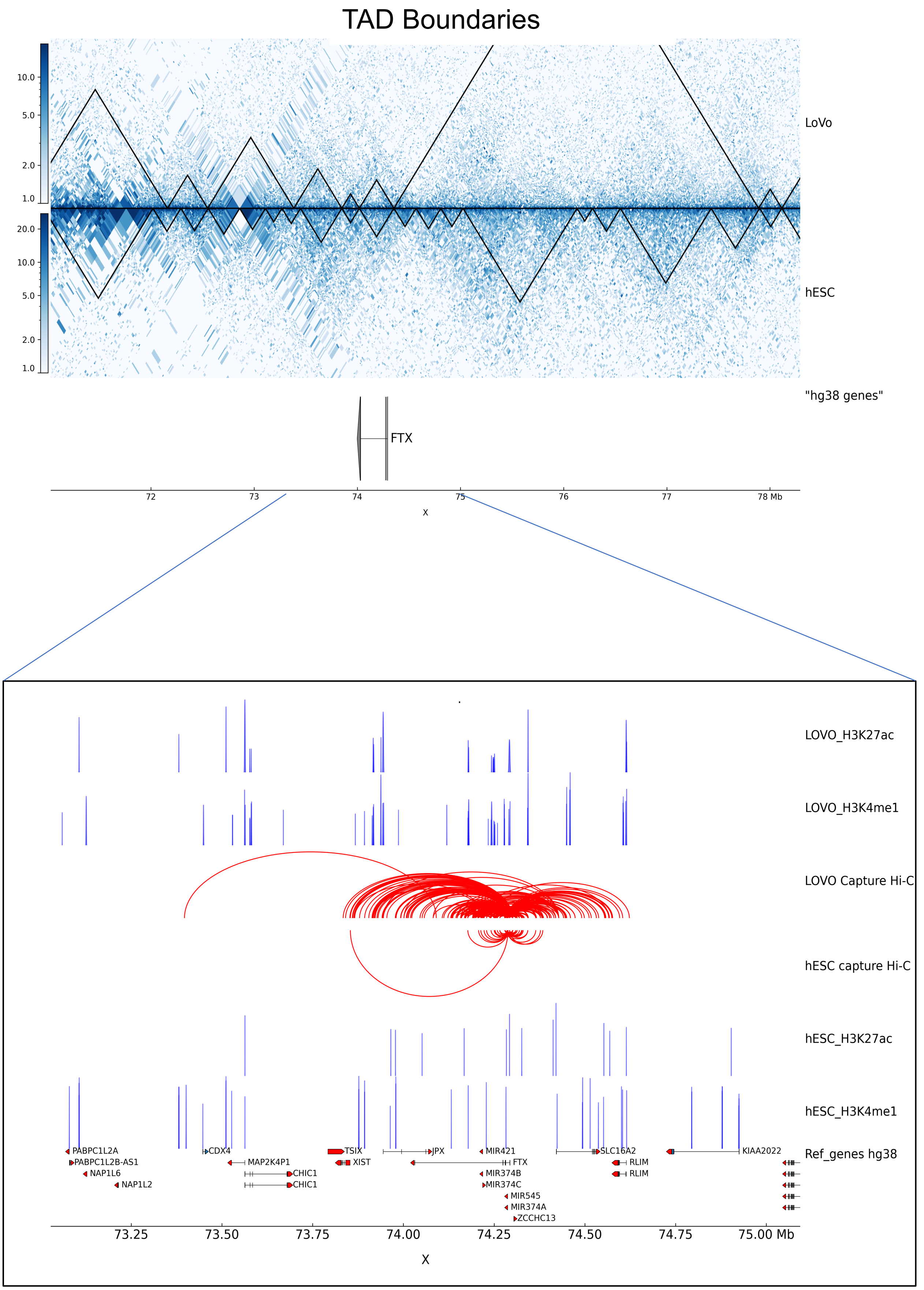

TAD Boundaries
LoVo
hESC
"hg38 genes"
FTX
72
73
74
75
76
77
78 Mb
X
LOVO_H3K27ac
LOVO_H3K4me1
LOVO Capture Hi-C
hESC capture Hi-C
hESC_H3K27ac
hESC_H3K4me1
Ref_genes hg38
PABPC1L2A
PABPC1L2B-AS1
NAP1L6
NAP1L2
CDX4
MAP2K4P1
CHIC1
CHIC1
TSIX
XIST
JPX
MIR421
FTX
MIR374B
MIR374C
MIR545
MIR374A
ZCCHC13
SLC16A2
RLIM
RLIM
KIAA2022
73.25
73.50
73.75
74.00
74.25
74.50
74.75
75.00 Mb
X

**Figure S6 (II). Effect on gene regulation due to structural changes between cancer (LoVo) versus normal (hESC) cell lines.** **(**A) ~ 7 Mb region of chromosome X encompassing the *FTX* gene is shown along with TADs boundaries of Hi-C interaction maps at 10 Kb resolution for case (LoVo) and control (hESC). (B) Zoomed-in view of the *FTX* locus in case (LoVo) and control (hESC) along with corresponding PCHi-C interaction, and ChIP-seq data for H3K27ac, H3K4me1 are displayed in blue peaks. Filtered FTX read counts used by CHiCAGO are displayed in red with the corresponding significant interactions shown as arcs. For clarity, only *FTX* interactions are shown.

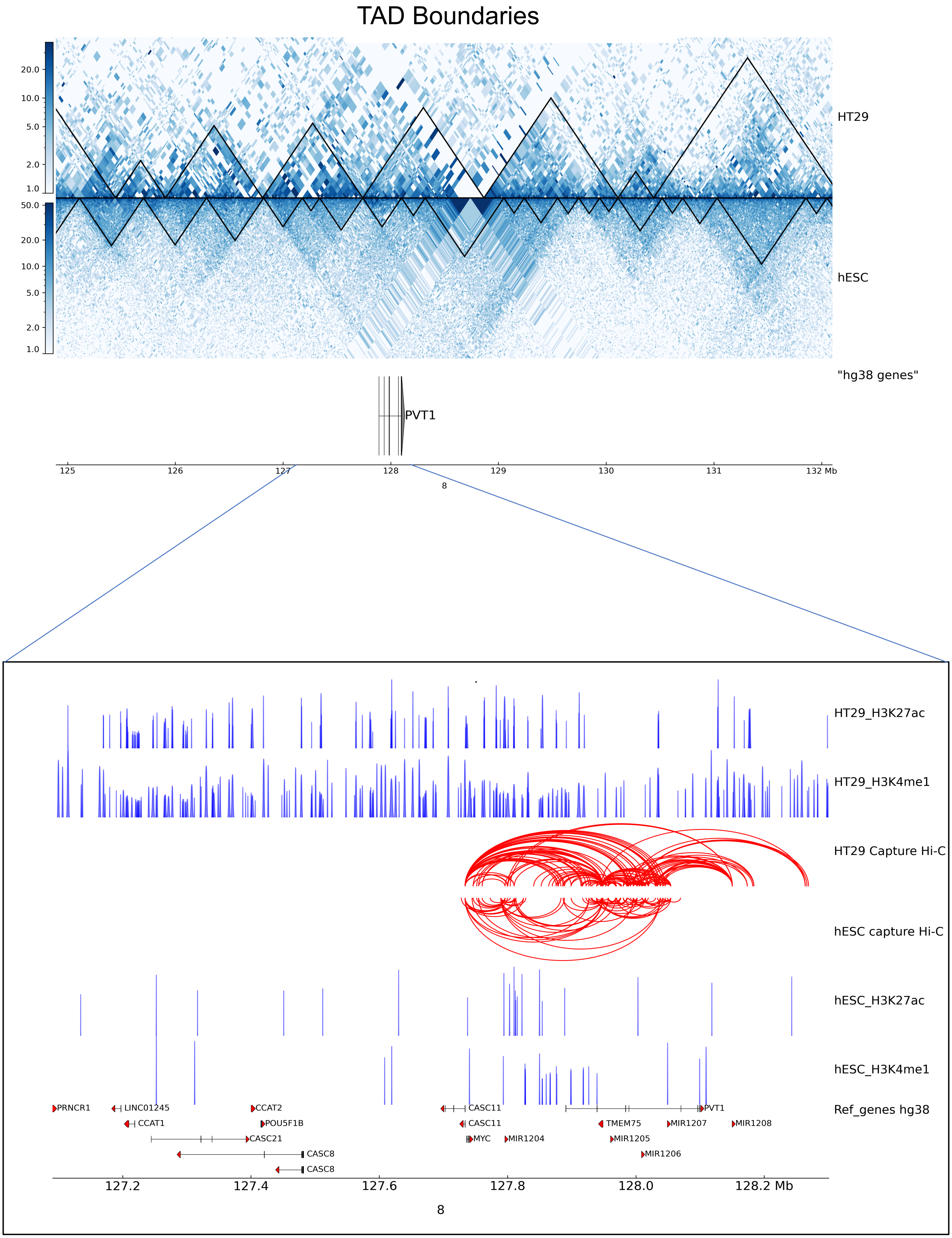

TAD Boundaries
HT29
hESC
"hg38 genes"
PVT1
125
126
127
128
129
130
131
132 Mb
8
HT29_H3K27ac
HT29_H3K4me1
HT29 Capture Hi-C
hESC capture Hi-C
hESC_H3K27ac
hESC_H3K4me1
Ref_genes hg38
PRNCR1
LINC01245
CCAT1
CCAT2
POU5F1B
CASC21
CASC8
CASC8
CASC11
CASC11
MYC
MIR1204
TMEM75
MIR1205
MIR1206
MIR1207
MIR1208
PVT1
127.2
127.4
127.6
127.8
128.0
128.2 Mb
8

**Figure S7 (I). Effect on gene regulation due to structural changes between cancer (HT29) versus normal (hESC) cell lines.** (A) ~ 7 Mb region of chromosome 8 encompassing the *PVT1* gene is shown along with TADs boundaries of Hi-C interaction maps at 10 Kb resolution for case (HT29) and control (hESC). (B) Zoomed-in view of the *PVT1* locus in case (HT29) and control (hESC) along with corresponding PCHi-C interaction, and ChIP-seq data for H3K27ac, H3K4me1 are displayed in blue peaks. Filtered PVT1 read counts used by CHiCAGO are displayed in red with the corresponding significant interactions shown as arcs. For clarity, only *PVT1* interactions are shown.

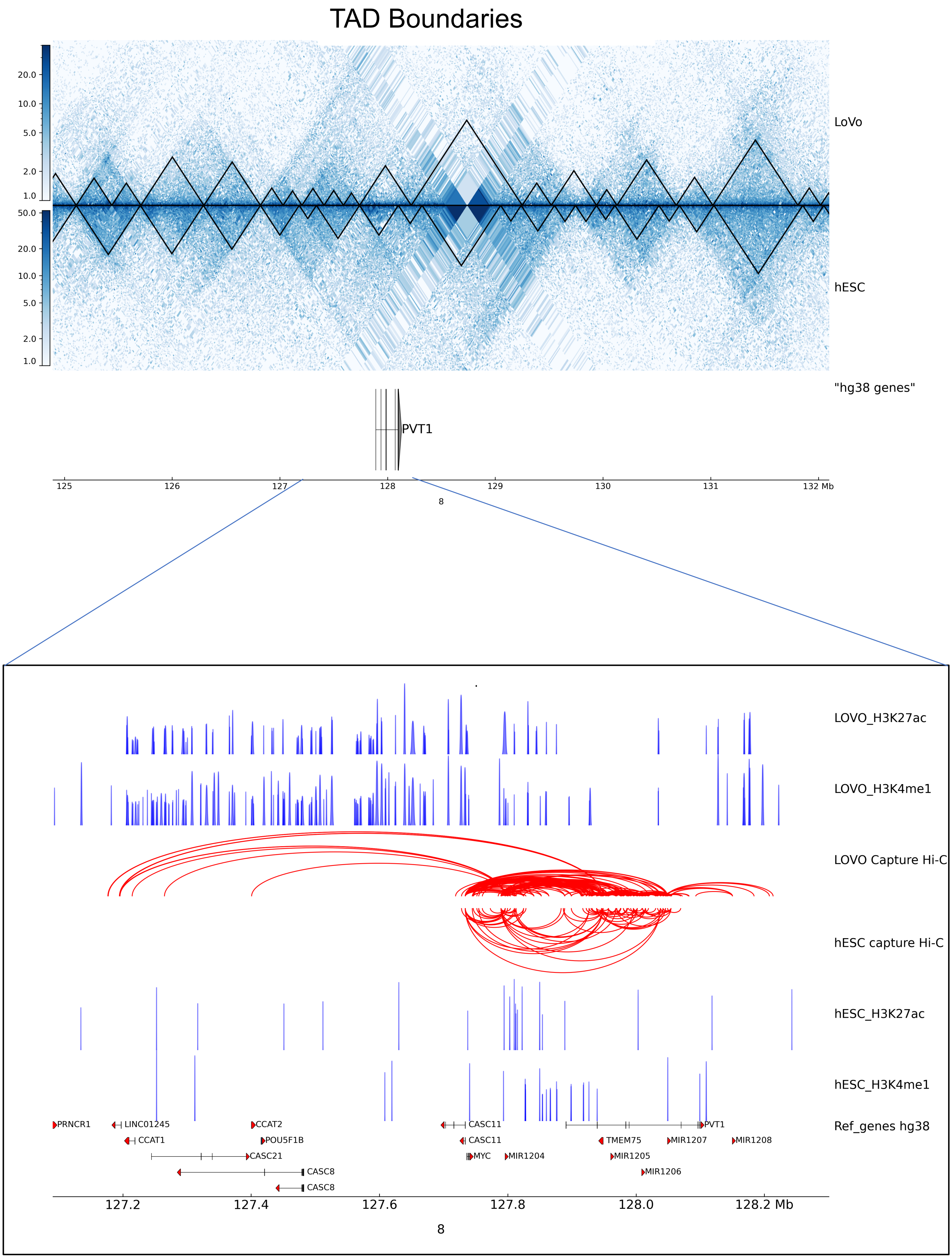

TAD Boundaries
LoVo
hESC
"hg38 genes"
PVT1
125
126
127
128
129
130
131
132 Mb
8
LOVO_H3K27ac
LOVO_H3K4me1
LOVO Capture Hi-C
hESC capture Hi-C
hESC_H3K27ac
hESC_H3K4me1
Ref_genes hg38
PRNCR1
LINC01245
CCAT1
CCAT2
POU5F1B
CASC21
CASC8
CASC8
CASC11
CASC11
MYC
MIR1204
TMEM75
MIR1205
MIR1206
MIR1207
MIR1208
PVT1
127.2
127.4
127.6
127.8
128.0
128.2 Mb
8

**Figure S7 (II). Effect on gene regulation due to structural changes between cancer (LoVo) versus normal (hESC) cell lines.** (A) ~ 7 Mb region of chromosome 8 encompassing the *PVT1* gene is shown along with TADs boundaries of Hi-C interaction maps at 10 Kb resolution for case (LoVo) and control (hESC). (B) Zoomed-in view of the PVT1 locus in case (LoVo) and control (hESC) along with corresponding PCHi-C interaction, and ChIP-seq data for H3K27ac, H3K4me1 are displayed in blue peaks. Filtered *PVT1* read counts used by CHiCAGO are displayed in red with the corresponding significant interactions shown as arcs. For clarity, only *PVT1* interactions are shown.

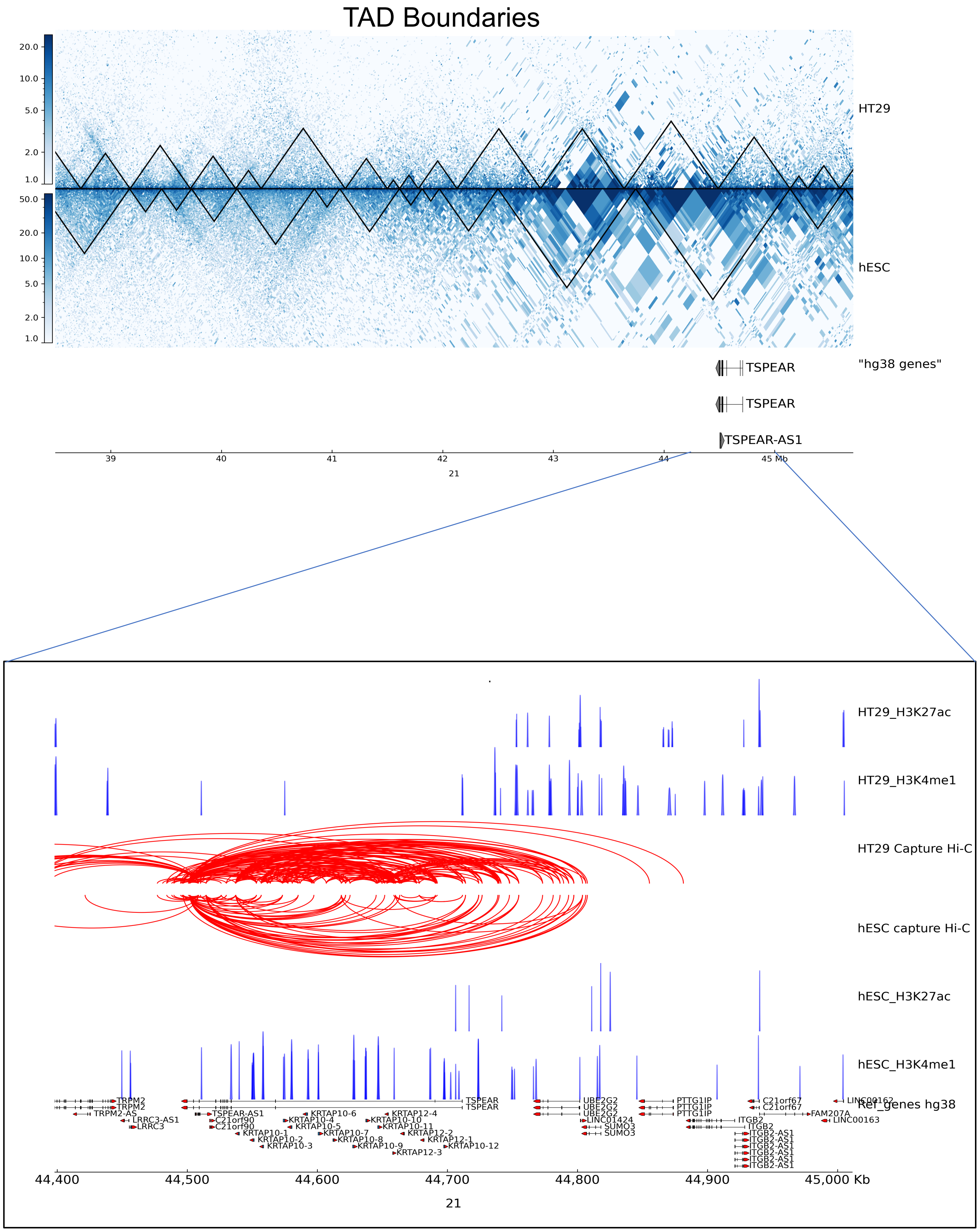
TAD Boundaries
HT29
hESC
"hg38 genes"
TSPEAR
TSPEAR
TSPEAR-AS1
39
40
41
42
43
44
45 Mb
21
HT29_H3K27ac
HT29_H3K4me1
HT29 Capture Hi-C
hESC capture Hi-C
hESC_H3K27ac
hESC_H3K4me1
Ref_genes hg38
44,400
44,500
44,600
44,700
44,800
44,900
45,000 Kb
21

**Figure S8 (I). Effect on gene regulation due to structural changes between cancer (HT29) versus normal (hESC) cell lines.** (A) ~ 7 Mb region of chromosome 21 encompassing the *TSPEAR* gene is shown along with TADs boundaries of Hi-C interaction maps at 10 Kb resolution for case (HT29) and control (hESC). (B) Zoomed-in view of the *TSPEAR* locus in case (HT29) and control (hESC) along with corresponding PCHi-C interaction, and ChIP-seq data for H3K27ac, H3K4me1 are displayed in blue peaks. Filtered TSPEAR read counts used by CHiCAGO are displayed in red with the corresponding significant interactions shown as arcs. For clarity, only *TSPEAR* interactions are shown.

### TAD Boundaries

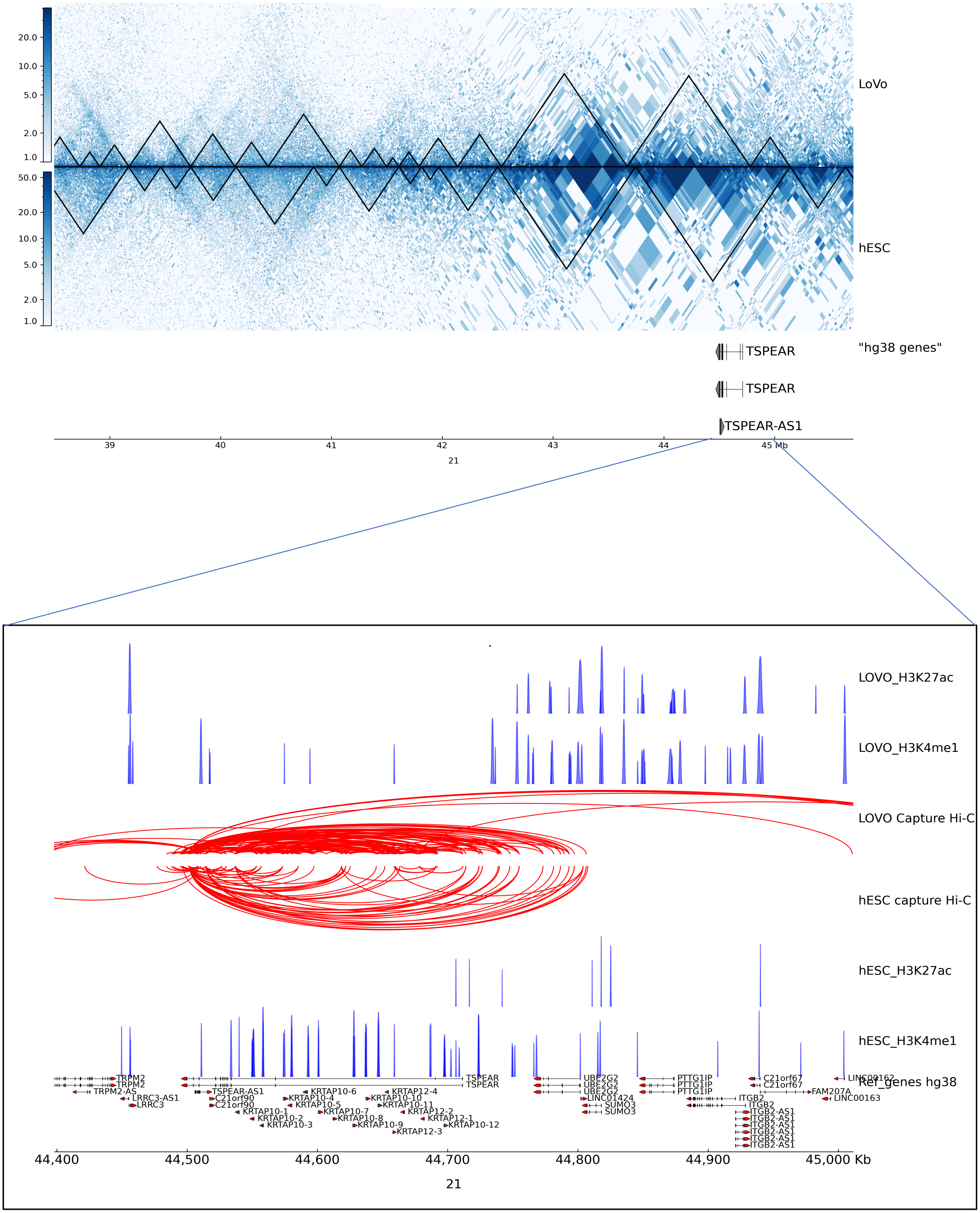

**Figure S8 (II). Effect on gene regulation due to structural changes between cancer (LoVo) versus normal (hESC) cell lines.** (A) ~ 7 Mb region of chromosome 21 encompassing the *TSPEAR* gene is shown along with TADs boundaries of Hi-C interaction maps at 10 kb resolution for case (LoVo) and control (hESC). (B) Zoomed-in view of the *TSPEAR* locus in case (LoVo) and control (hESC) along with corresponding PCHi-C interaction, and ChIP-seq data for H3K27ac, H3K4me1 are displayed in blue peaks. Filtered TSPEAR read counts used by CHiCAGO are displayed in red with the corresponding significant interactions shown as arcs. For clarity, only *TSPEAR* interactions are shown.

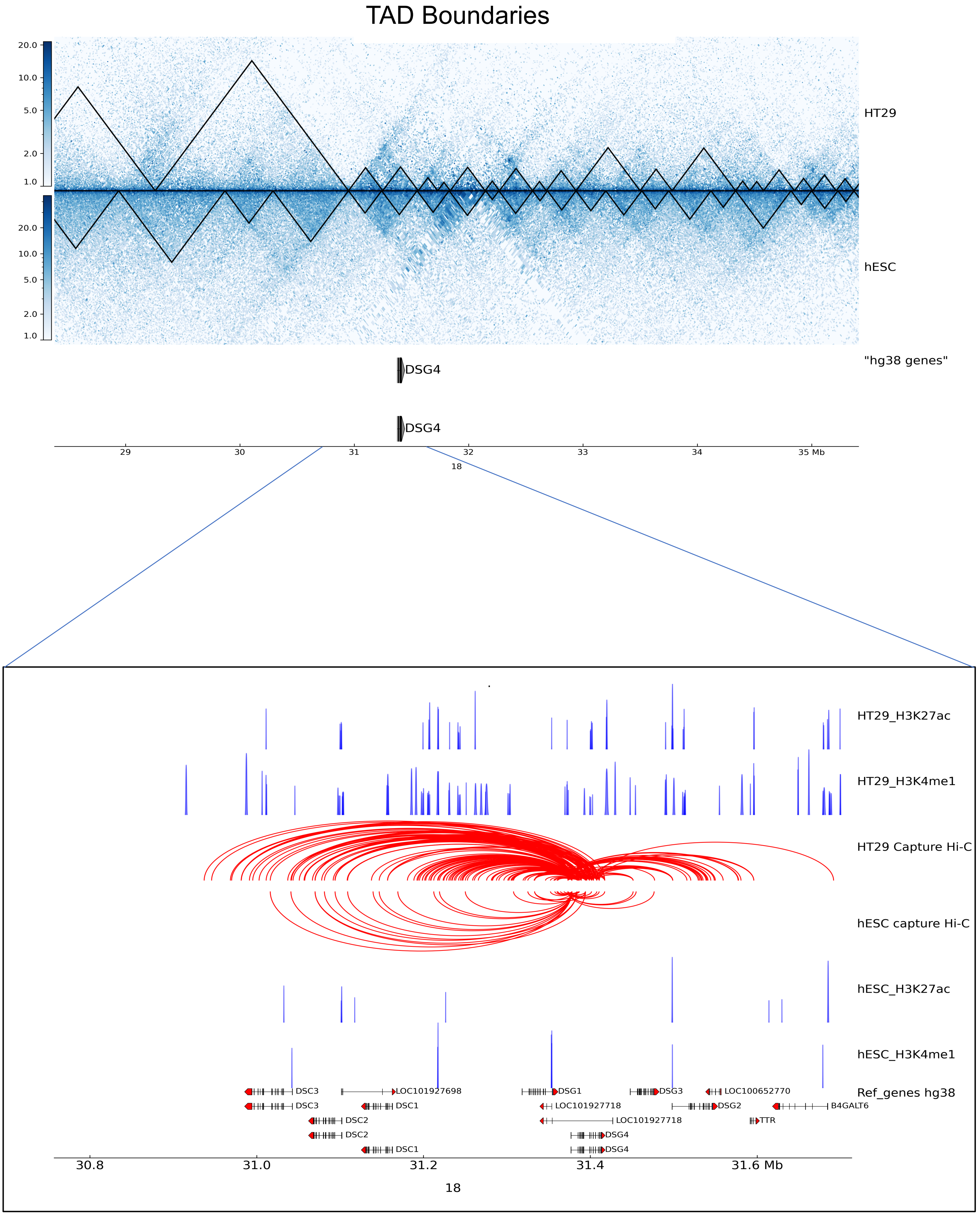
TAD Boundaries
HT29
hESC
"hg38 genes"
DSG4
DSG4
29
30
31
32
33
34
35 Mb
18
HT29_H3K27ac
HT29_H3K4me1
HT29 Capture Hi-C
hESC capture Hi-C
hESC_H3K27ac
hESC_H3K4me1
Ref_genes hg38
DSC3
LOC101927698
DSG1
DSG3
LOC100652770
DSC1
LOC101927718
DSG2
B4GALT6
DSC2
TTR
DSG4
30.8
31.0
31.2
31.4
31.6 Mb
18

**Figure S9 (I). Effect on gene regulation due to structural changes between cancer (HT29) versus normal (hESC) cell lines.** (A) ~ 7 Mb region of chromosome 18 encompassing the *DSG4* gene is shown along with TADs boundaries of Hi-C interaction maps at 10 Kb resolution for case (HT29) and control (hESC). (B) Zoomed-in view of the *DSG4* locus in case (HT29) and control (hESC) along with corresponding PCHi-C interaction, and ChIP-seq data for H3K27ac, H3K4me1 are displayed in blue peaks. Filtered DSG4 read counts used by CHiCAGO are displayed in red with the corresponding significant interactions shown as arcs. For clarity, only *DSG4* interactions are shown.

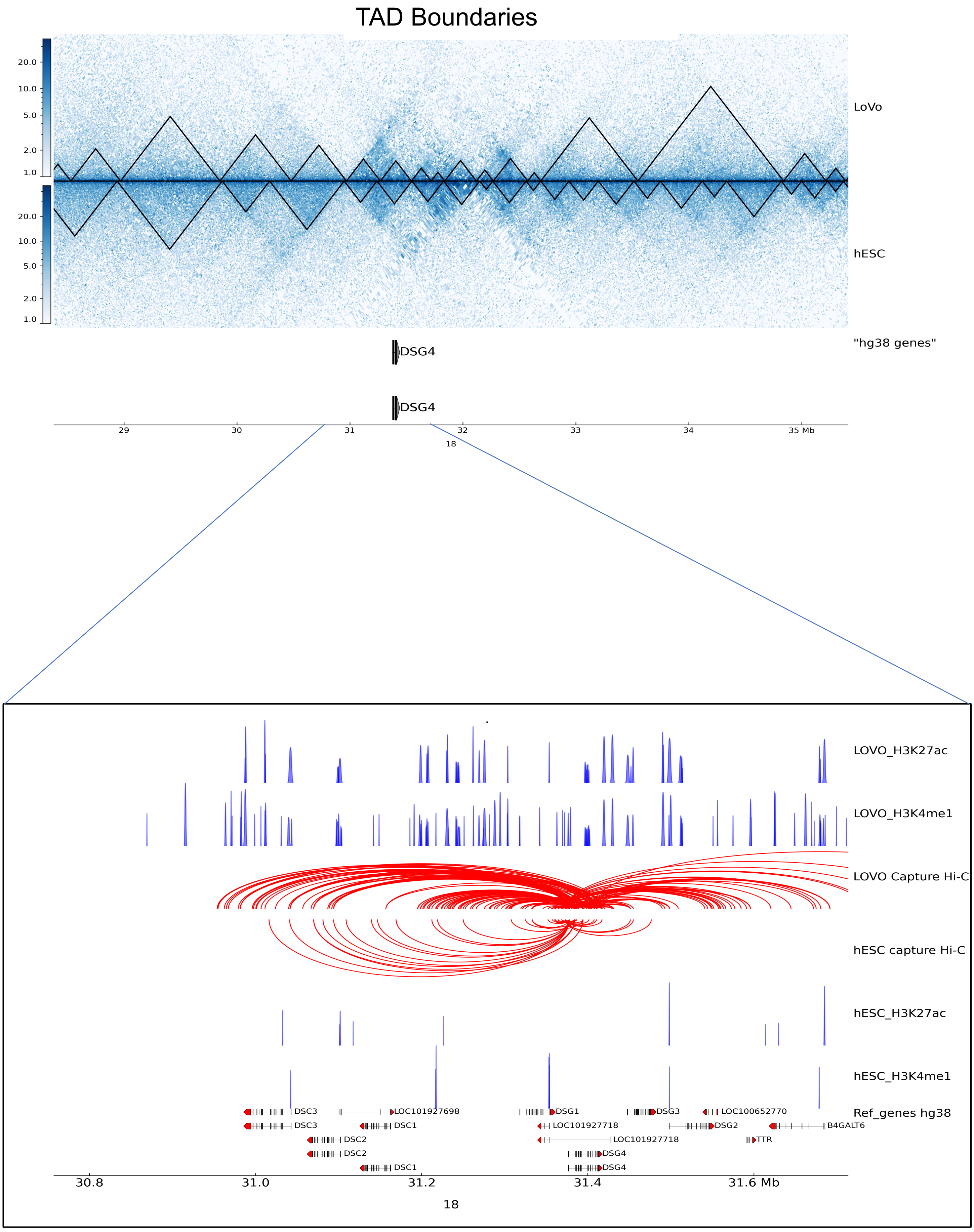
TAD Boundaries
LoVo
hESC
"hg38 genes"
DSG4
DSG4
29
30
31
32
33
34
35 Mb
18
LOVO_H3K27ac
LOVO_H3K4me1
LOVO Capture Hi-C
hESC capture Hi-C
hESC_H3K27ac
hESC_H3K4me1
Ref_genes hg38
DSC3
DSC3
LOC101927698
DSC1
DSC2
DSC2
DSC1
DSG1
LOC101927718
LOC101927718
DSG4
DSG4
DSG3
DSG2
LOC100652770
B4GALT6
TTR
30.8
31.0
31.2
31.4
31.6 Mb
18

**Figure S9 (II). Effect on gene regulation due to structural changes between cancer (LoVo) versus normal (hESC) cell lines.** (A) ~ 7 Mb region of chromosome 18 encompassing the *DSG4* gene is shown along with TADs boundaries of Hi-C interaction maps at 10 Kb resolution for case (LoVo) and control (hESC). (B) Zoomed-in view of the DSG4 locus in case (LoVo) and control (hESC) along with corresponding PCHi-C interaction, and ChIP-seq data for H3K27ac, H3K4me1 are displayed in blue peaks. Filtered DSG4 read counts used by CHiCAGO are displayed in red with the corresponding significant interactions shown as arcs. For clarity, only *DSG4* interactions are shown.

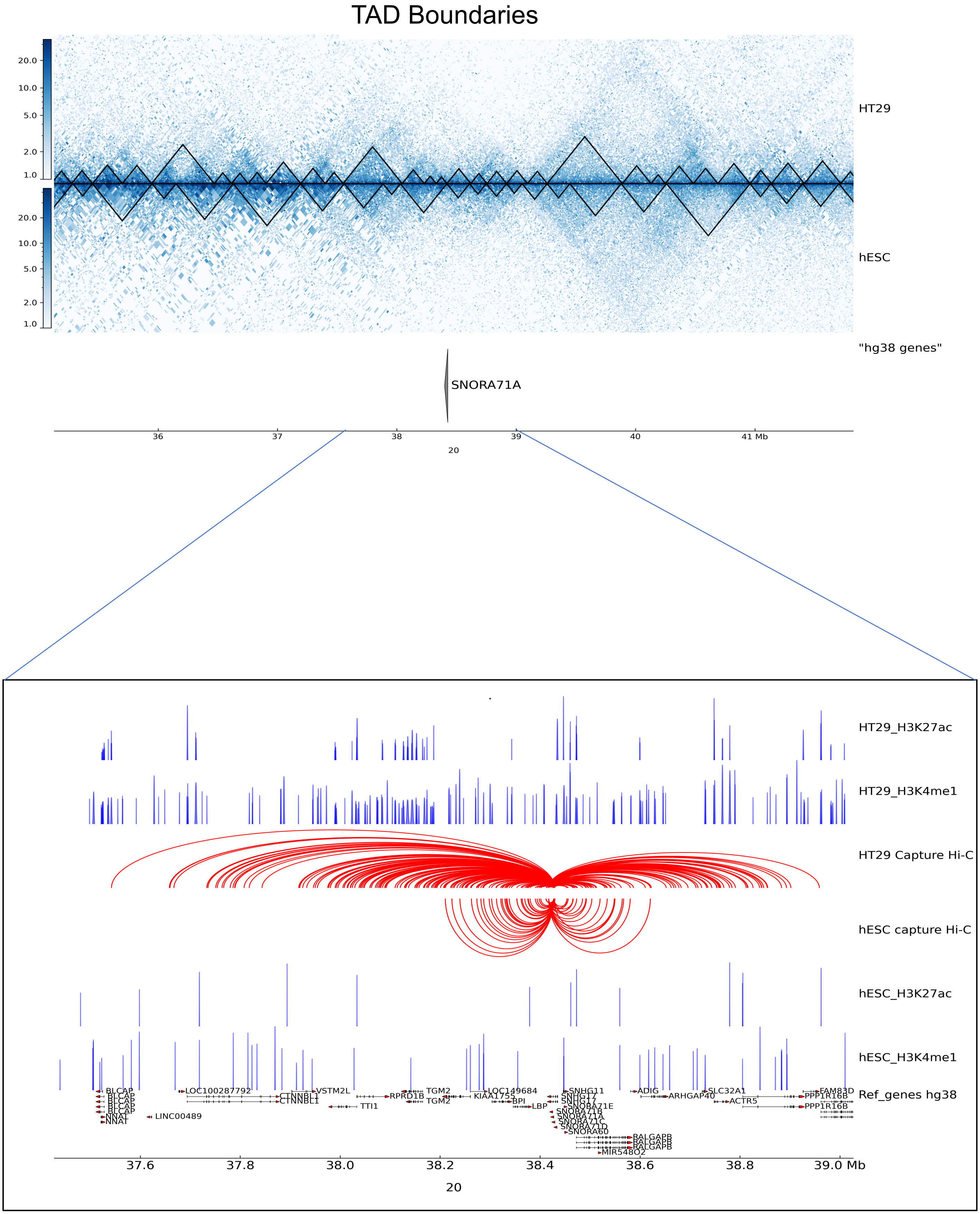

TAD Boundaries
HT29
hESC
"hg38 genes"
SNORA71A
36
37
38
39
40
41 Mb
20
HT29_H3K27ac
HT29_H3K4me1
HT29 Capture Hi-C
hESC capture Hi-C
hESC_H3K27ac
hESC_H3K4me1
Ref_genes hg38
BLCAP
NNAT
LINC00489
LOC100287792
CTNNBL1
VSTM2L
TTI1
RPRD1B
TGM2
KIAA1755
LOC149684
BPI
LBP
SNHG11
SNHG17
SNORA71B
SNORA71E
SNORA71A
SNORA71C
SNORA71D
SNORA60
RALGAPB
MIR548O2
ADIG
ARHGAP40
SLC32A1
ACTR5
PPP1R16B
FAM83D
37.6
37.8
38.0
38.2
38.4
38.6
38.8
39.0 Mb
20

**Figure S10 (I). Effect on gene regulation due to structural changes between cancer (HT29) versus normal (hESC) cell lines. (**A) ~ 6 Mb region of chromosome 20 encompassing the *SNORA71A* gene is shown along with TADs boundaries of Hi-C interaction maps at 10 Kb resolution for case (HT29) and control (hESC). (B) Zoomed-in view of the SNORA71A locus in case (HT29) and control (hESC) along with corresponding PCHi-C interaction, and ChIP-seq data for H3K27ac, H3K4me1 are displayed in blue peaks. Filtered SNORA71A read counts used by CHiCAGO are displayed in red with the corresponding significant interactions shown as arcs. For clarity, only *SNORA71A* interactions are shown.

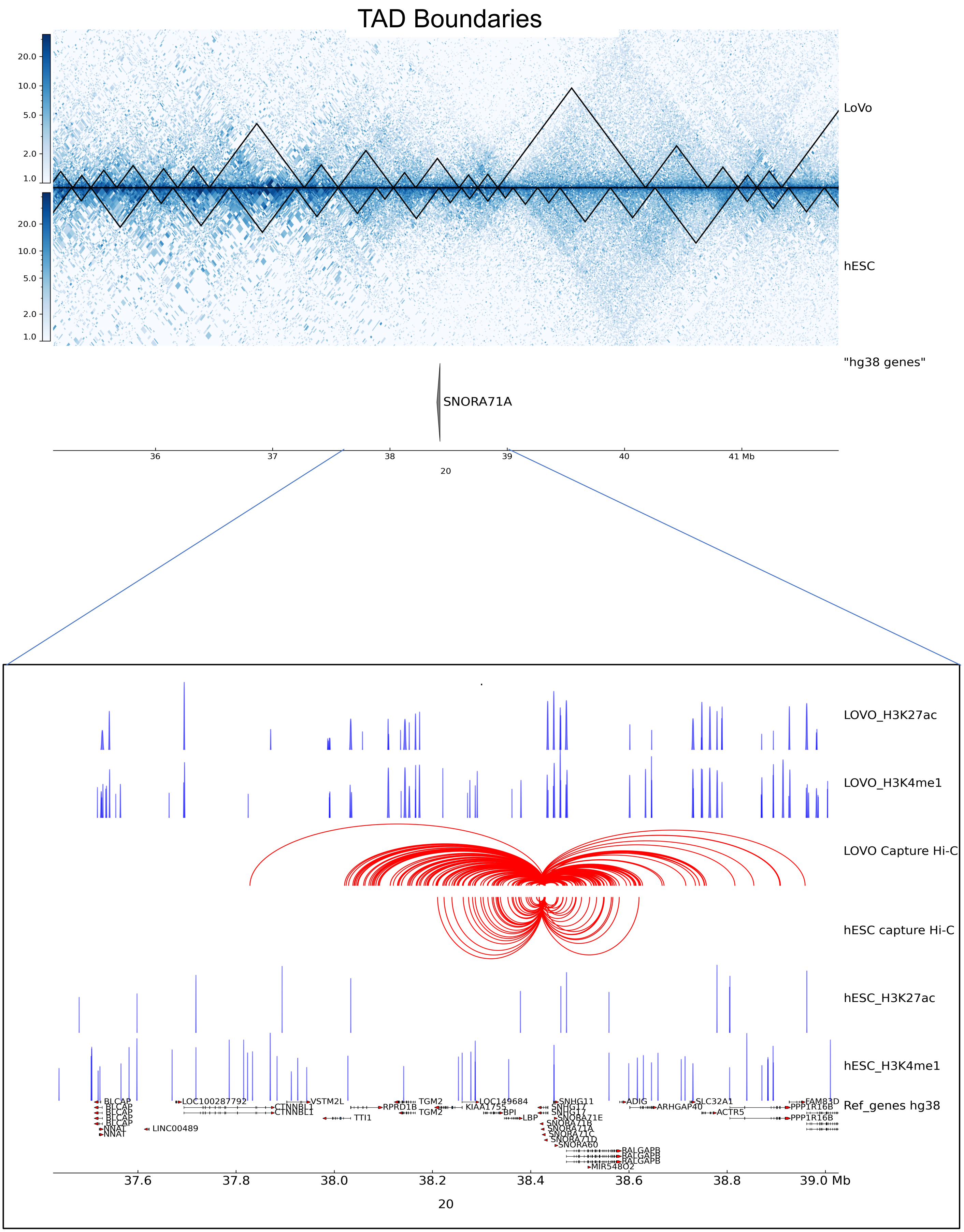

TAD Boundaries
LoVo
hESC
"hg38 genes"
SNORA71A
36
37
38
39
40
41 Mb
20
LOVO_H3K27ac
LOVO_H3K4me1
LOVO Capture Hi-C
hESC capture Hi-C
hESC_H3K27ac
hESC_H3K4me1
Ref_genes hg38
BLCAP
NNAT
LINC00489
LOC100287792
CTNNBL1
VSTM2L
TTI1
RPRD1B
TGM2
KIAA1755
LOC149684
BPI
LBP
SNHG11
SNHG17
SNORA71E
SNORA71B
SNORA71A
SNORA71C
SNORA71D
SNORA60
RALGAPB
MIR548O2
ADIG
ARHGAP40
SLC32A1
ACTR5
FAM83D
PPP1R16B
37.6
37.8
38.0
38.2
38.4
38.6
38.8
39.0 Mb
20

**Figure S10 (II). Effect on gene regulation due to structural changes between cancer (LoVo) versus normal (hESC) cell lines.** (A) ~ 6 Mb region of chromosome 20 encompassing the *SNORA71A* gene is shown along with TADs boundaries of Hi-C interaction maps at 10 Kb resolution for case (LoVo) and control (hESC). (B) Zoomed-in view of the *SNORA71A* locus in case (LoVo) and control (hESC) along with corresponding PCHi-C interaction, and ChIP-seq data for H3K27ac, H3K4me1 are displayed in blue peaks. Filtered SNORA71A read counts used by CHiCAGO are displayed in red with the corresponding significant interactions shown as arcs. For clarity, only *SNORA71A* interactions are shown.

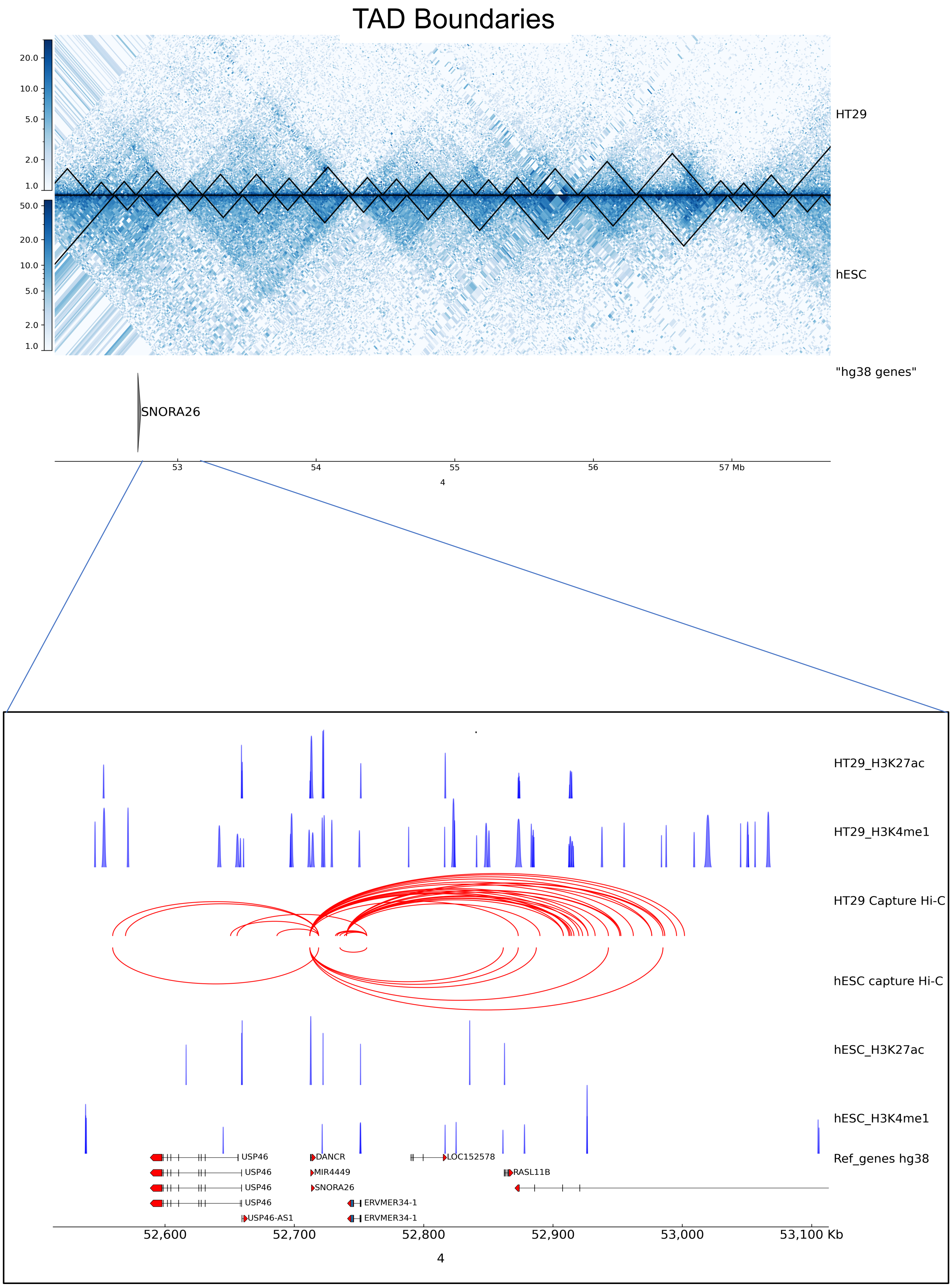
TAD Boundaries
HT29
hESC
"hg38 genes"
SNORA26
53
54
55
56
57 Mb
4
HT29_H3K27ac
HT29_H3K4me1
HT29 Capture Hi-C
hESC capture Hi-C
hESC_H3K27ac
hESC_H3K4me1
Ref_genes hg38
USP46
USP46
USP46
USP46
USP46-AS1
DANCR
MIR4449
SNORA26
ERVMER34-1
ERVMER34-1
LOC152578
RASL11B
52,600
52,700
52,800
52,900
53,000
53,100 Kb
4

**Figure S11 (I). Effect on gene regulation due to structural changes between cancer (HT29) versus normal (hESC) cell lines.** (A) ~ 5 Mb region of chromosome 4 encompassing the *SNORA26* gene is shown along with TADs boundaries of Hi-C interaction maps at 10 Kb resolution for case (HT29) and control (hESC). (B) Zoomed-in view of the *SNORA26* locus in case (HT29) and control (hESC) along with corresponding PCHi-C interaction, and ChIP-seq data for H3K27ac, H3K4me1 are displayed in blue peaks. Filtered SNORA26 read counts used by CHiCAGO are displayed in red with the corresponding significant interactions shown as arcs. For clarity, only *SNORA26* interactions are shown.

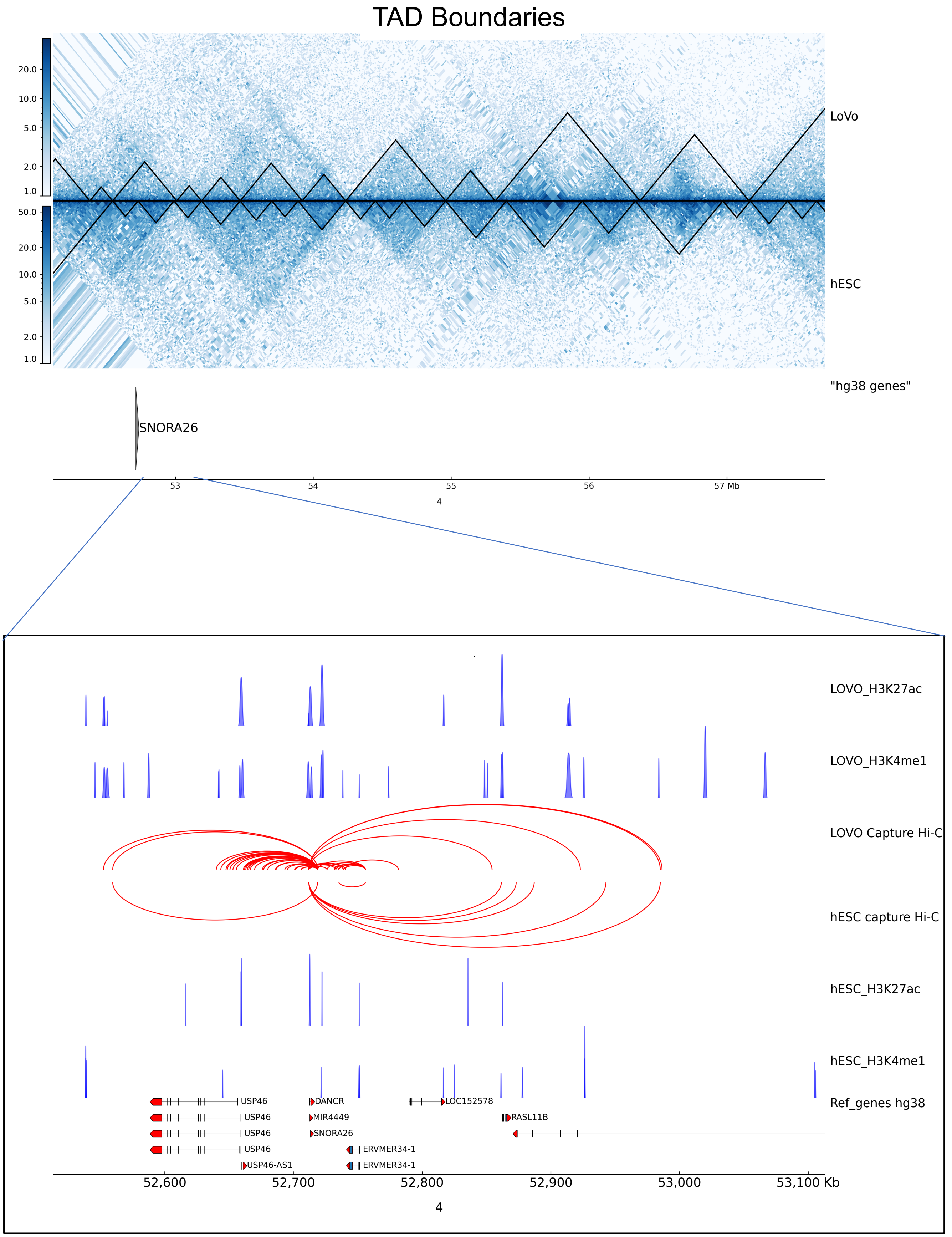
TAD Boundaries
20.0
10.0
5.0
2.0
1.0
50.0
20.0
10.0
5.0
2.0
1.0
LoVo
hESC
"hg38 genes"
SNORA26
53
54
55
56
57 Mb
4
LOVO_H3K27ac
LOVO_H3K4me1
LOVO Capture Hi-C
hESC capture Hi-C
hESC_H3K27ac
hESC_H3K4me1
Ref_genes hg38
USP46
USP46
USP46
USP46
USP46-AS1
DANCR
MIR4449
SNORA26
ERVMER34-1
ERVMER34-1
LOC152578
RASL11B
52,600
52,700
52,800
52,900
53,000
53,100 Kb
4

**Figure S11 (II). Effect on gene regulation due to structural changes between cancer (LoVo) versus normal (hESC) cell lines.** (A) ~ 5 Mb region of chromosome 4 encompassing the *SNORA26* gene is shown along with TADs boundaries of Hi-C interaction maps at 10 Kb resolution for case (LoVo) and control (hESC). (B) Zoomed-in view of the *SNORA26* locus in case (LoVo) and control (hESC) along with corresponding PCHi-C interaction, and ChIP-seq data for H3K27ac, H3K4me1 are displayed in blue peaks. Filtered SNORA26 read counts used by CHiCAGO are displayed in red with the corresponding significant interactions shown as arcs. For clarity, only *SNORA26* interactions are shown.
