## Supplementary material for "Combined promoter-capture Hi-C and Hi-C analysis reveals a fine-tuned regulation of 3D chromatin architecture in colorectal cancer": Saw et al., Supplementary_table_list

**Supplementary Table S1.** List of oligonucleotide primer sequences used for qRT-PCR analysis.

**Supplementary Table S2**. List of primer sequences used for ChIP-PCR analysis.

**Supplementary Table S3.** List of the transcripts along with the gene annotations. This list has been generated using TCGA colorectal cancer transcript database.

**Supplementary Table S4.** List contains enriched gene ontology and pathways for both HT29 and LoVo cell lines.

**Supplementary Table S5.** The number of overlaps of bait’s interactions (region of interest) with regulatory elements such as promoter-like (PLS), proximal enhancer-like (pELS), distal enhancer-like (dELS), DNase-H3K4me3 and CTCF regions which were downloaded from the ENCODE database (https://screen-v2.wenglab.org).
