## Supplementary_table_S5 for "Combined promoter-capture Hi-C and Hi-C analysis reveals a fine-tuned regulation of 3D chromatin architecture in colorectal cancer"

Supplementary file

(bait interaction with regulatory elements stage-wise)

| **Bait interaction with ↓** | **hESC/HT29** | | | | | | **hESC/LOVO** | | | | | |
|---|---|---|---|---|---|---|---|---|---|---|---|---|
| | hESC | | | HT29 | | | hESC | | | LOVO | | |
| | I | II | III | I | II | III | I | II | III | I | II | III |
| Promoter region | 20521 | 33115 | 26967 | 44715 | 65586 | 52929 | 20494 | 33063 | 21144 | 43437 | 65357 | 37467 |
| Proximal enhancer region | 77329 | 106326 | 83454 | 199317 | 243287 | 187422 | 76196 | 105711 | 75822 | 183535 | 225550 | 152083 |
| Distal enhancer region | 238403 | 153654 | 111216 | 926554 | 550096 | 395886 | 230179 | 146125 | 97088 | 812700 | 477218 | 318545 |
| Dnase-H3K4me3 region | 5934 | 4259 | 3177 | 21104 | 14163 | 10788 | 5718 | 3933 | 2257 | 18695 | 12831 | 6246 |
| CTCF region | 11391 | 5766 | 3900 | 48654 | 23452 | 16213 | 11091 | 5430 | 3620 | 41875 | 19951 | 13256 |

Table E1: Total number of bait interaction with regulatory elements stage-wise (I, II and III).

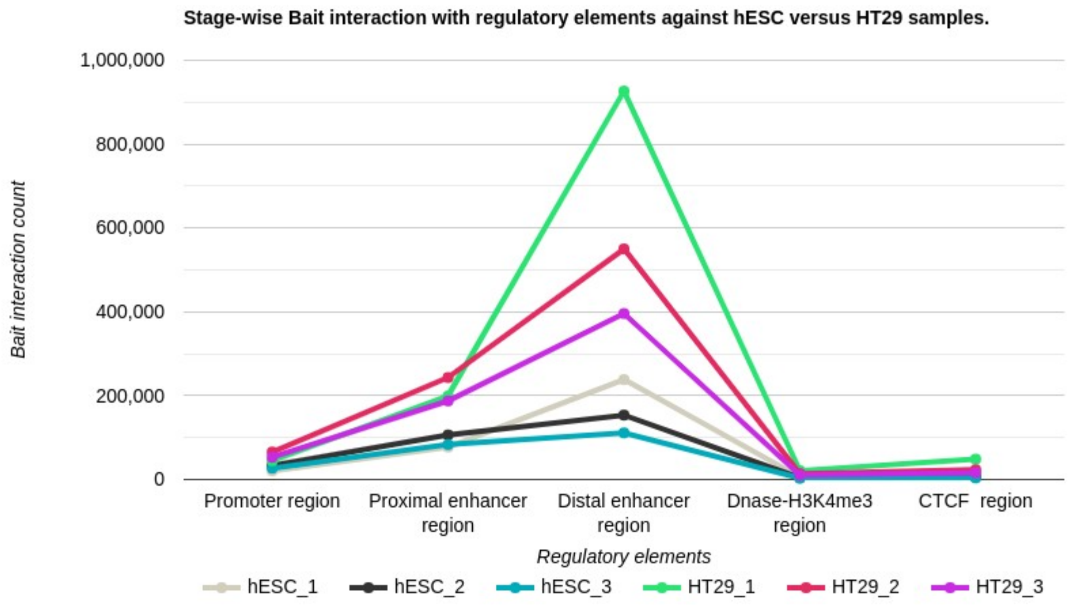


Figure E2: Stage-wise total number of bait interaction with regulatory elements (Promoter-like (PLS), Proximal enhancer-like (pELS), Distal enhancer-like (dELS), DNase-H3K4me3 and CTCF regions) hESC and HT29 cell lines in hESC/HT29.

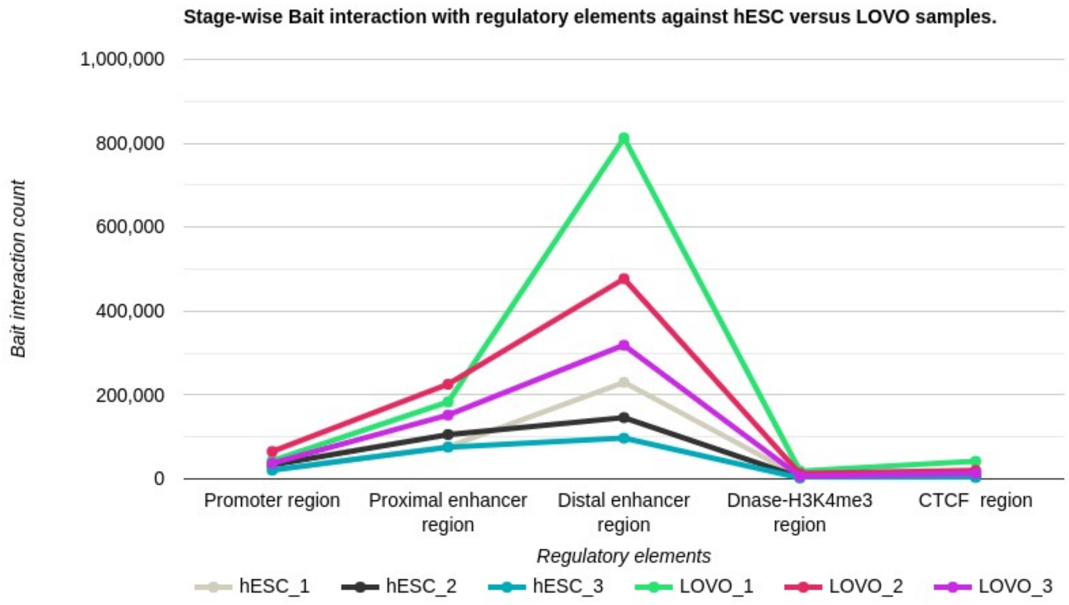


Figure E3: Stage-wise total number of bait interaction with regulatory elements (Promoter-like (PLS), Proximal enhancer-like (pELS), Distal enhancer-like (dELS), DNase-H3K4me3 and CTCF regions) hESC and LOVO cell lines in hESC/LOVO.
